# Escape from replicative crisis and immortalization precedes complex genome evolution

**DOI:** 10.64898/2026.10.09.758087

**Authors:** Tobias T. Schmidt, Marco R. Cosenza, Tessa M. Popay, Sara Przetocka, Candy Haggblom, Nicholas Allsing, Kathryn Lande, Julia W. Grimstead, April E. Williams-Wehner, Duncan M. Baird, Jesse R. Dixon, Jan O. Korbel, Jan Karlseder

## Abstract

Shortening of telomeres restricts the proliferative capacity of human somatic cells by inducing replicative senescence and crisis. During cancerous transformation, premalignant cells must overcome these barriers and acquire replicative immortality. Single-cell DNA strand-specific genome sequencing of crisis cells revealed extreme genome instability characterized by complex chromosomal abnormalities. Unexpectedly, spontaneously emerging postcrisis clones showed relatively few and simple structural variants. However, with accumulating divisions following crisis escape, the genomes of immortal postcrisis clones evolve and amass genomic alterations, including tetraploidization events with subsequent losses and gains of individual chromosomes. Our data point towards a new model of tumorigenesis in which crisis acts as a strong selective barrier against cells with aberrant karyotypes, and in which high karyotypic complexity is acquired after immortalization and crisis escape.

## Main Text

Genome instability is a major driver of evolution and frequently the cause or the defining characteristic of human pathologies including cancer. In the case of cancer, genomic instability not only facilitates malignant transformation by promoting the acquisition of cancer hallmarks (*1*) but also contributes to intra-tumor heterogeneity and ongoing evolution of cancers following transformation (*2*). Particularly, high chromosomal instability (CIN) is linked to the most aggressive, recurrent and metastatic cancers (*3–7*). One essential hallmark of cancer and a fundamental difference between cancer and human somatic cells is replicative immortality (*1*). Cancer cells acquire replicative immortality by activating a telomere maintenance mechanism (TMM) that counteracts telomere shortening. While the majority of cancers reactivate the reverse transcriptase telomerase (*8*), 10-15% of cancers maintain their telomeres with the recombination-based Alternative Lengthening of Telomeres (ALT) mechanism (*9*).

Telomeres are the protective, nucleoprotein structures at the end of eukaryotic linear chromosomes. In human somatic cells, telomeres shorten with every round of DNA replication due to the “end-replication problem” and subsequent processing (*10–12*). Short telomeres are recognized by the DNA damage response, marked as telomere dysfunction-induced foci (TIF) and trigger a p53 and retinoblastoma (Rb) checkpoint-dependent permanent cell cycle arrest called replicative senescence (*13*). Inactivation of DNA damage checkpoints by either genetic or epigenetic alterations or expression of viral oncogenes such as the human papilloma virus (HPV) E6 and E7 proteins or the simian virus 40 large T (SV40LT) antigen, facilitates senescence bypass and extends proliferation with continuous telomere shortening until cells with critically short, deprotected telomeres encounter a second telomere-dependent proliferation barrier called replicative crisis (*14–18*). Crisis in fibroblasts and epithelial cells is characterized by extensive telomere fusions, widespread genomic instability and autophagy-dependent, innate immunity-driven cell death (*17–21*). Very infrequently, one in 10-30 million fibroblast cells will spontaneously escape crisis by activating a TMM or through another unknown mechanism (*22–24*). Thus, replicative crisis represents a stringent barrier against human malignant transformation that premalignant cells must overcome during cancer initiation.

Genetically engineered mouse models have yielded fundamental insights into oncogenic drivers and tumor suppressors. However, they are less well suited to investigate early human tumorigenesis, particularly the acquisition of cellular immortality and the genome evolution that occurs during cancer initiation (*25*). This limitation stems from two fundamental species differences: First, mouse cells express telomerase in most tissues (*26*) leading to stable telomere length over their lifespan (*27*), and second, laboratory mice have extremely long telomeres (*28*) that can sustain the production of viable offspring without overt pathology for multiple generations, even in the absence of any TMM (*29*). In contrast, human somatic cells lack telomerase (*30*), leading to telomere shortening and crisis (*14–18*). Consequently, the early stages of human cancer initiation remain incompletely understood.

Here, we utilize a well-established, spontaneous in vitro crisis escape and immortalization model to investigate human cancer initiation. We found that crisis is characterized by high CIN, however, cells which escape crisis and acquire replicative immortality show surprisingly stable karyotypes with only simple structural variants (SVs), suggesting that complex chromosomal alterations are strongly counter-selected in the absence of a TMM. However, the immortal postcrisis clones, that driven by an altered 3D organization of the telomerase (TERT) locus maintain their telomeres by telomerase, continued to evolve acquiring more complex karyotypes. Our results therefore support a model in which the highly rearranged genomes characteristic of some cancers (*7, 31*) are generated primarily after crisis escape and the acquisition of replicative immortality.

## Results

### Short telomeres drive complex chromosomal abnormalities in crisis

To model replicative crisis in vitro, we expressed the HPV E6 and E7 oncogenes in IMR90 human lung fibroblasts (IMR90 ^E6E7^) and grew them to crisis. As telomere attrition-dependent genome instability is the underlying trigger for cell death during replicative crisis, we first analyzed telomere length by nanopore telomere sequencing (Telo-seq) (*32*) of growing (population doubling (PD) 61) and crisis (PD105) IMR90 ^E6E7^ cells. In crisis, telomeres are shorter relative to growing cells (median 2225 base pairs (bp) vs. 4740 bp), the fraction of telomeres shorter than one kilobase (kb) accumulates (15.45% vs. 1.71%) and telomeres become more heterogeneous in length (coefficient of variation (CV) 0.74 vs. 0.53) (Fig. 1A, and table S1). Next, using a custom IMR90 ^E6E7^ reference genome (see Materials and Methods), we analyzed chromosome arm-specific telomere length at 40 of the 46 chromosome arms (Fig. 1B, fig. S1A-C, and Data S1). At every analyzed chromosome arm the mean telomere length was shorter in crisis relative to growing IMR90 ^E6E7^ cells (fig. S1D and Data S1) and average telomere shortening varied between 22 and 125 bp per PD at different chromosome arms (fig. S1E and Data S1). While the intrasample mean chromosome arm-specific telomere length were heterogeneous and varied up to 5.7 and 5.9 kb in growing and crisis IMR90 ^E6E7^ cells, respectively (Fig. 1B, fig. S1C and Data S1), the relative order of chromosome arms in both samples was conserved (fig. S1F).

**Fig. 1.**
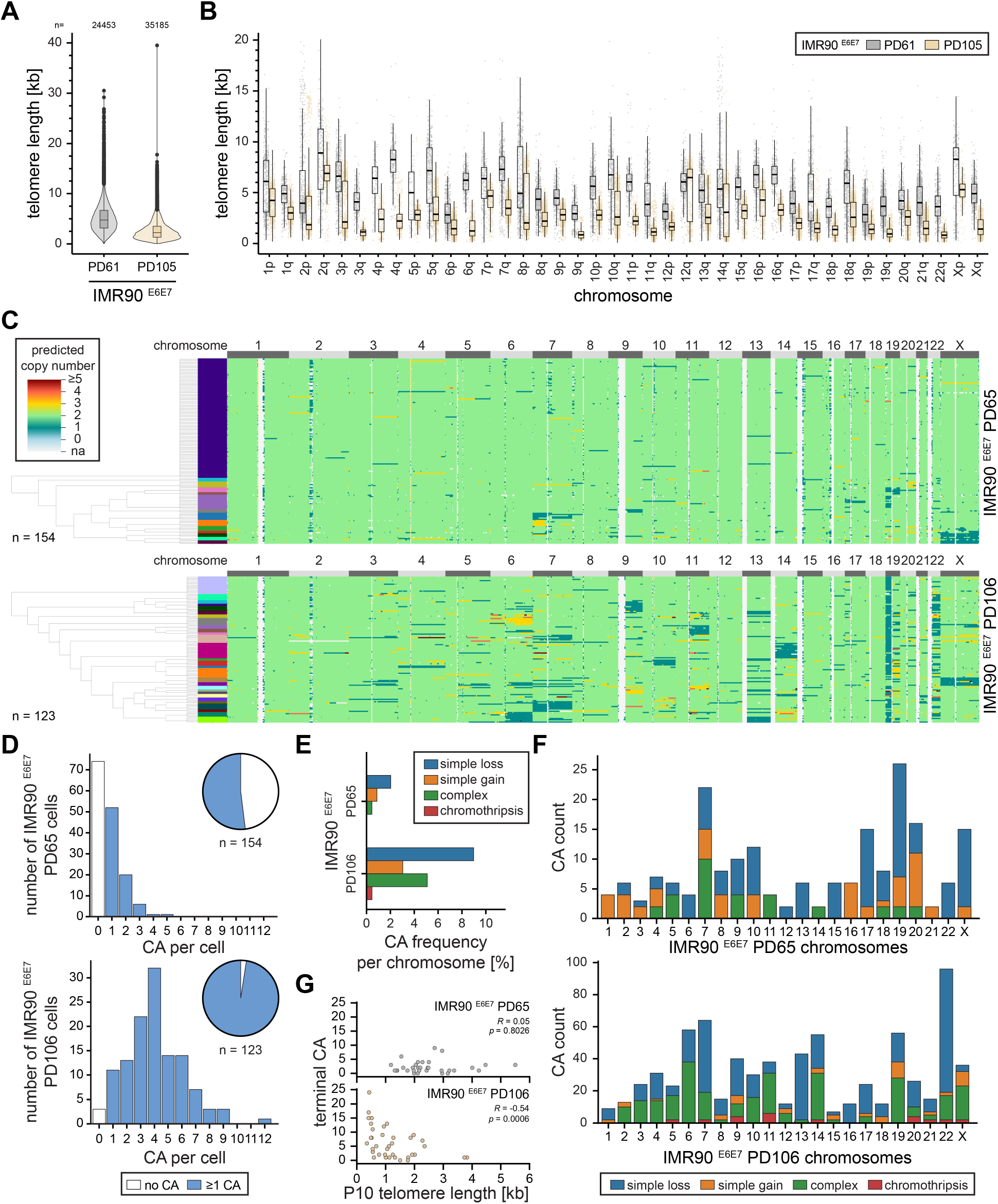
Crisis cells experience high heterogeneous genome instability. (**A**) Violin plot of Telo-seq telomere length measurements in IMR90 ^E6E7^ growing (population doubling (PD) 61) and crisis (PD105) cells. Violin represents telomere length distribution in kilobases (kb). Boxplot shows the median telomere length with interquartile range (IQR) and whiskers represent 1.5-fold IQR. The number of telomeric reads per sample is shown above the plot. One experiment. For more detail see table S1. (**B**) Boxplot of chromosome arm-specific telomere length in kb of growing and crisis IMR90 ^E6E7^ cells based on mapping against a custom IMR90 ^E6E7^ reference genome with mapping quality 40 and higher. The middle line represents median, the box the IQR and whiskers the 1.5-fold IQR. Individual telomere reads are shown as points. The acrocentric p-arms of chromosomes 13, 14, 15, 21, and 22 and chromosome arm 20p could not be assembled in the custom reference genome and thus, are not shown. One experiment. For more detail see Data S1. (**C**) Copy number estimates of individual IMR90 ^E6E7^ growing (PD65) and crisis (PD106) cells based on single-cell Strand-seq. Phylogenetic trees were calculated with PICASSO and the most representative tree of eight PICASSO runs is shown. (**D**) Histogram of number of chromosomal abnormalities (CA) per cell in IMR90 ^E6E7^ growing (PD65) and crisis (PD106) cells. (**E**) Bar graph of CA frequency per chromosome in percent in growing (PD65) and crisis (PD106) IMR90 ^E6E7^ cells. (**F**) Bar graph of CAs per chromosome in growing (PD65) and crisis (PD106) IMR90 ^E6E7^ cells. (**G**) Scatter plot of the number of terminal CA against the 10^th^ percentile (P10) telomere length in kb for growing and crisis IMR90 ^E6E7^ cells. Spearman’s rank correlation was performed.

Consistent with the short telomeres detected in crisis and previous reports (*17–21*), IMR90 ^E6E7^ crisis cells showed elevated numbers of 53BP1 foci, a marker of DNA damage, increased metaphasic TIFs, fusions and micronuclei (fig. S2A-E). Given the telomere length heterogeneity (Fig. 1A,B, fig. S1C, table S1 and Data S1), we hypothesized that the spontaneous telomere dysfunction-driven mutational events in crisis are also heterogeneous and therefore required single-cell resolution. Thus, we performed single-cell strand-specific genome sequencing (Strand-seq) followed by computational analysis with strandtools (*33–35*) to comprehensively resolve chromosomal abnormalities (CAs) in individual growing and crisis cells. Growing, young IMR90 ^E6E7^ (PD65) cells had on average one distinctive CA per cell, and in 48.1% (74 of 154) of cells no CA was detectable (Fig. 1C,D and fig. S2A). In contrast, IMR90 ^E6E7^ crisis cells (PD106) carried an average of four CAs per cell and only 2.4% (three of 123) cells had no detectable CA (Fig. 1C,D and fig. S2F). While the CA spectra in growing IMR90 ^E6E7^ cells were mainly dominated by simple losses, some gains and few complex CAs (defined as copy number changes with more than one breakpoint), crisis IMR90 ^E6E7^ cells were characterized by a high CA burden together with an increase in more complex CAs, that were frequently involving terminal alterations (Fig. 1E and fig. S2G). The complex CAs observed in crisis included CA patterns indicative of breakage-fusion-bridge (BFB) cycles (*36*) (fig. S2G,H) as well as chromothripsis, a mutational event in which genetic material is shattered and randomly assembled (*37, 38*) (Fig. 1E, and fig. S2H), and in one instance a highly amplified region indicative of extrachromosomal DNA (fig. S2H). In IMR90 ^E6E7^ crisis cells, all chromosomes and arms carried CAs (Fig. 1F and fig. S2I). Complex CAs were detected at 20 different chromosomes whereas chromothripsis was found at 9 different chromosomes (Fig. 1F and fig. S2I). Similar results were obtained with a second IMR90 ^E6E7^ replicate independently transduced with HPV E6 and E7 oncogenes and crisis WI38 human lung fibroblasts expressing the SV40LT antigen (WI38 ^SV40LT^), which both showed an increased heterogeneous mutational burden during crisis (fig. S3). Finally, to address whether the short telomeres in IMR90 ^E6E7^ crisis cells (Fig. 1A,B, Data S1) drive CAs in crisis, we compared chromosome arm-specific terminal CAs and the 10^th^ percentile (P10) of telomere length with Spearman’s rank correlation coefficient analysis. In growing IMR90 ^E6E7^ cells terminal CAs did not correlate with the P10 telomere length, however, terminal CAs were significantly increased at crisis on chromosome arms with the shortest telomeres (Spearman’s R = −0.54, *p* = 0.0006, Fig. 1G). Thus, crisis cells experience a high and heterogeneous genome instability burden with complex CAs, which is driven by short telomeres.

### Spontaneous crisis escape is accompanied by relatively few and simple structural variants

Replicative crisis represents a stringent barrier against human malignant transformation (*22*). However, it is plausible that the high genome instability in crisis (Fig. 1 and fig. S2) may facilitate crisis escape and the acquisition of cancer hallmarks (*1*). To model the early stages of malignant transformation in vitro, we isolated spontaneous postcrisis clones from a large population of IMR90 ^E6E7^ crisis cells. For this, we grew IMR90 ^E6E7^ cells to crisis, seeded 81 million crisis cells at low density and cultured them for 60 days. At this time, the majority of cells had died in crisis, and spontaneously emerging postcrisis clones were isolated and expanded (Fig. 2A). Growth curve analysis of eight postcrisis clones revealed three distinct growth behaviors (Fig. 2B) (please also see Discussion and Limitation of the study sections): First, postcrisis clones PC1, 6, 45 and 48 grew initially, but then entered a growth plateau and died, suggesting that these mortal postcrisis clones passed crisis with no or insufficient TMM. Second, similar to the mortal postcrisis clones, the postcrisis clone PC7 entered a growth plateau marked by widespread cell death but subsequently emerged from a 50-day long plateau and continued to grow for at least 250 days. This indicates that PC7 was initially isolated as a mortal clone (PC7early) with no or insufficient TMM but acquired immortality during the ensuing growth plateau following crisis (PC7late). Third, the postcrisis clones PC37, 39 and 42 grew linearly for more than 300 days indicating that these postcrisis clones escaped crisis with sufficient TMM activity to support growth and were therefore considered as immortal.

**Fig. 2.**
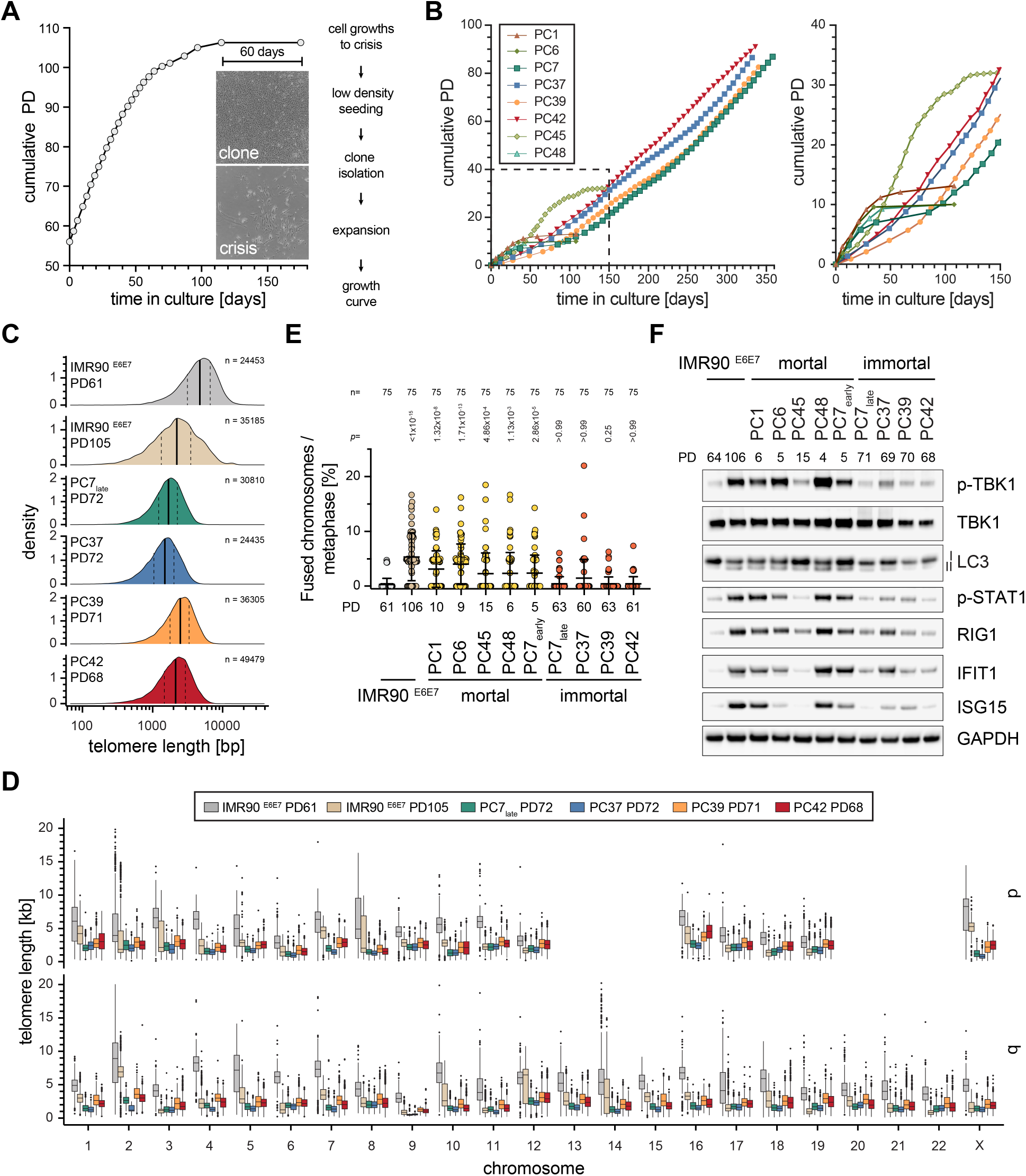
Spontaneously emerging immortal postcrisis clones have short telomeres and an intermediate DNA damage and innate immune signature. (**A**) Spontaneous IMR90 ^E6E7^ postcrisis clone isolation scheme. Precrisis growth of IMR90 ^E6E7^ is shown as cumulative population doublings (PDs) against days in culture. (**B**) Growth curve analysis of isolated postcrisis clones. Right plot represents the first 150 days in culture as shown in the dashed black line in the left plot. (**C**) Density plot of log10 transformed telomere length distribution in base pairs (bp) measured by Telo-seq in indicated cells. Vertical solid line represents median telomere length with the 25% and 75% quantiles shown as vertical dotted lines. PD61 and PD105 are growing and crisis IMR90 ^E6E7^ cells, respectively, and are displayed as controls and have been shown in Fig. 1A. One experiment. For more detail see table S1. (**D**) Boxplot of chromosome arm-specific telomere length in kilobase (kb) of growing and crisis IMR90 ^E6E7^ cells and immortal postcrisis clones based on mapping against a custom IMR90 ^E6E7^ reference genome with mapping quality 40 and higher. The middle line represents median, the box the interquartile range (IQR) and whiskers the 1.5-fold IQR and outliers are shown as black points. The acrocentric p-arms of chromosomes 13, 14, 15, 21, and 22 and chromosome arm 20p could not be assembled in the custom reference genome and thus, are not shown. One experiment. Growing and crisis IMR90 ^E6E7^ cells are shown as reference and previously reported in Fig. 1B. For more details see Data S1. (**E**) Scatter dot plot of the percentage of fused chromosomes per metaphase with mean and standard deviation. Each dot represents one metaphase. Cumulative data of three independent experiments with 25 metaphase each. Statistical test: Kruskal-Wallis test with Dunn’s multiple comparison correction compared to IMR90 ^E6E7^ PD61 growing control. IMR90 ^E6E7^ PD61 and PD106 are also shown in fig. S2C. (**F**) Western blot analysis of indicated samples. IMR90 ^E6E7^ PD64 and PD106 represent growing and crisis controls, respectively. Three independent experiments.

In the isolated postcrisis clones, bulk whole genome sequencing revealed a consensus karyotype with large-scale gains and losses, while highly complex SVs patterns where largely absent (fig. S4A). Overall, the immortal postcrisis clones acquired about twice the amount of SVs compared to mortal postcrisis clones (fig. S4B) and somatic molecular clock mutational signatures showed enrichment in all postcrisis clones (fig. S4C) (*39*). All postcrisis clones exhibited losses of chromosome arms 13q and 22q (fig. S4A), consistent either with their origin from the same crisis cell subpopulation or with convergent evolution. Notably, both alterations were also found in a subpopulation of individual crisis cells, and subclonal architecture analysis indicated that they had been acquired independently, in line with convergent evolution. Furthermore, all postcrisis clones except PC42 showed lesions at chromosome 19 (fig. S4A). In crisis, both arms of chromosome 19 are among the ten arms with the shortest telomeres (Fig. 1B, Data S1) and chromosome 19 is frequently altered in individual crisis cells (Fig. 1C). Interestingly, three of the four immortal postcrisis clones acquired a subtetraploid genome (fig. S4A). Taken together, these results show that the isolated postcrisis clones escaped crisis with relatively few and simple SVs, suggesting a strong negative selection against highly complex SVs during crisis. At the same time, changes affecting chromosomes 13, 19 and 22 are recurrent in postcrisis clones, with independent acquisition of chromosome losses and frequent involvement of chromosomes harboring the shortest telomeres.

### Immortal postcrisis clones exhibit short telomeres, ongoing genome instability and an intermediate type I interferon response

Consistent with a higher replicative age (Fig. 2A,B and fig. S4C), all mortal and immortal postcrisis clones except PC39 had shorter or equal telomere length than crisis cells, as determined by Single Telomere Length Analysis (STELA) (*40*) (fig. S5A,B) and Telo-seq (*32*) (Fig. 2C,D, fig. S5C,D, table S1 and Data S1). Bulk and chromosome arm-specific telomere length were more homogenous in comparison to growing and crisis cells (Fig. 2C,D, table S1, and Data S1). Further, chromosome arm-specific telomere length analysis revealed that the TMM active in the immortal postcrisis clones preferentially stabilizes or elongates the telomeres at the chromosome arms with the shortest telomeres in crisis. Conversely, the arms with relatively long telomeres in crisis are several kb shorter in the immortal postcrisis clones (Fig. 2D and fig. S5E,F). Thus, in immortal postcrisis clones TMM stabilizes telomeres, but they do remain relatively short compared to pre-crisis lengths.

Next, we analyzed markers of genome instability in the mortal and immortal postcrisis clones in comparison to IMR90 ^E6E7^ growing and crisis cells. In accordance with their inability to acquire sufficient telomere maintenance, the mortal postcrisis clones display heterogeneous, yet elevated markers of genome instability (Fig. 2E and fig. S6). Chromosome fusions were largely reduced in the immortal postcrisis clones (Fig. 2E and fig. S6D), whereas metaphasic TIFs, 53BP1 foci and micronuclei show intermediate values between those of growing and crisis IMR90 ^E6E7^ cells (fig. S6A-C). Taken together, despite having solved the end-replication problem, immortal postcrisis cells continue to experience genome instability after crisis escape.

Crisis cells experience autophagic cell death and cGAS-STING-TBK1 innate immune signaling with an interferon-stimulated gene (ISG) signature (*17, 18*), Consistent with this, we detected increased levels of LC3II, a marker of autophagy, phosphorylation of TBK1 and STAT1 at serine 172 and threonine 701, respectively, as well as upregulation of three ISGs (IFIT1, ISG15 and RIG1) in IMR90 ^E6E7^ crisis relative to growing control cells using western blotting (Fig. 2F). LC3II levels remained elevated in all mortal clones but were reduced in the immortal postcrisis clones. In the mortal postcrisis clones the markers of innate immune signaling resembled IMR90 ^E6E7^ crisis cells except for clone PC45 that had reduced levels. In the immortal postcrisis clones, the innate immune signaling was reduced relative to crisis, but elevated compared to growing IMR90 ^E6E7^ controls. Differential expression analysis of the crisis and postcrisis bulk RNA transcriptomes relative to the IMR90 ^E6E7^ growing controls revealed that in crisis and postcrisis cells the response to type I interferon was upregulated (fig. S7A) (*17*). In addition, the ISG signature found in crisis relative to growing IMR90 ^E6E7^ controls (fig. S7B) (*17*) was significantly upregulated in postcrisis cells relative to growing cells, but downregulated relative to crisis cells (fig. S7C,D). Of note, immortal PC7late cells showed reduced innate immune response signaling relative to the mortal PC7early cells sampled prior to the plateau (Fig. 2B,F and fig. S7).

Next, we performed soft-agar assays to investigate whether the isolated immortal postcrisis clones acquired the ability to grow anchorage-independent during crisis escape, a feature of transformed cells (*41*). Whereas HeLa LT cancer cells formed detectable colonies in soft agar, all immortal postcrisis clones and the parental IMR90 ^E6E7^ control were incapable of anchorage-independent growth (fig. S8A).

Finally, we analyzed the upregulated differentially expressed genes of the immortal postcrisis clones relative to both growing and crisis IMR90 ^E6E7^ cells to inquire about the possible acquisition of oncogenic characteristics. Interestingly, all immortal postcrisis clones overexpress the epidermal growth factor receptor (EGFR), a frequent oncogene in cancer (*42*)(fig. S8B-D). Further, EGFR protein levels were similar between different PDs of immortal postcrisis clones suggesting that EGFR overexpression is not a consequence of selection during postcrisis growth but rather acquired during crisis and crisis escape (fig. S8E).

Taken together, the immortal postcrisis clones have short telomeres, experience ongoing genome instability and have an intermediate innate immune response relative to growing and crisis IMR90 ^E6E7^, possibly related to ongoing genome instability. While the immortal postcrisis cells are not yet fully transformed, the EGFR upregulation in the immortal postcrisis clones exemplifies that oncogenes can be upregulated during crisis and crisis escape.

### Immortal postcrisis clones maintain their telomeres by epigenetic activation of telomerase

To evaluate how the immortal postcrisis clones maintain their telomeres, we measured telomerase activity using Telomeric Repeat Amplification Protocol (TRAP) (*43*). Whereas no telomerase activity was detectable in the parental IMR90 ^E6E7^ and mortal PC45 postcrisis cells, telomerase activity was detectable in all four immortal postcrisis clones (Fig. 3A). Further, the expression of the protein component of telomerase TERT, measured by reverse transcriptase quantitative PCR (RT-qPCR), was increased in the immortal postcrisis clones, but not in the mortal PC45 and PC7early postcrisis cells (fig. S9A). The latter suggests that indeed the postcrisis clone PC7 initially escaped crisis with insufficient TMM and upregulated TERT during the 50-day growth plateau to acquire full replicative immortality. Taken together, all four immortal postcrisis clones maintain their telomeres by reactivation of telomerase.

**Fig. 3.**
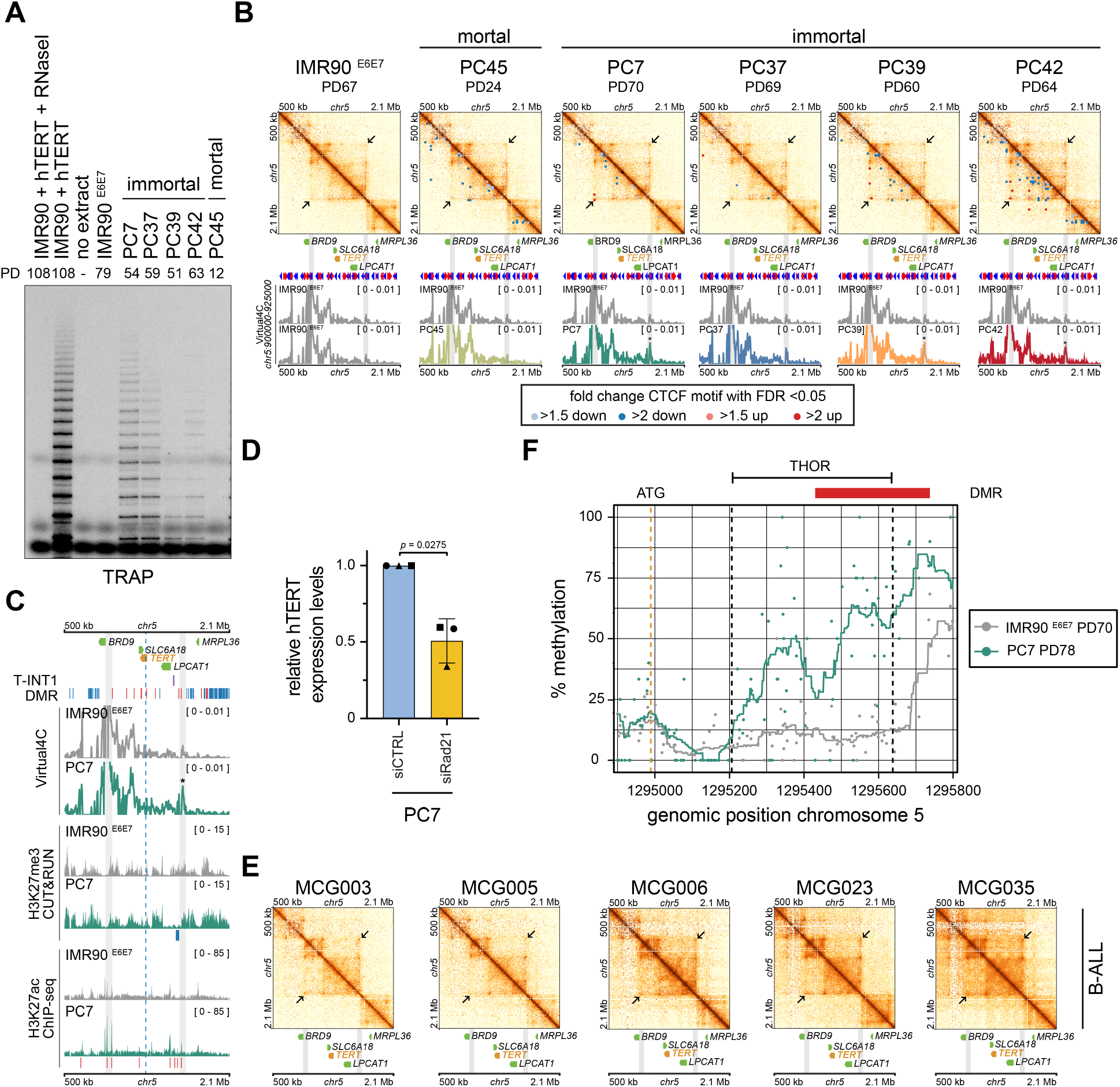
Altered genome organization promotes TERT expression. (**A**) Telomerase Repeat Amplification Protocol (TRAP) of parental IMR90 ^E6E7^ and postcrisis clones. RNaseI treated or untreated extracts of IMR90 cells ectopically overexpressing hTERT are used as negative and positive control, respectively. Representative of two independent experiments. (**B**) Hi-C sequencing analysis of IMR90 ^E6E7^ and postcrisis clones. The surrounding of the TERT locus (chr5:500000-2100000, GRCh38/hg38) is shown as contact map with Virtual4C (chr5:900000-925000, GRCh38/hg38). The detected loop anchor is marked with an arrow and if statistically significant (FDR < 0.05) with an asterisk in Virtual4C. Differential CTCF contacts are color coded according to fold change. Two replicates per sample. (**C**) H3K27me3, H3K27ac and DNA methylation in IMR90 ^E6E7^ (PD64-72) and PC7 (PD76-82) were measured by Cut&Run, ChIP-seq and Oxford nanopore technologies (ONT) sequencing, respectively. Differential peaks (FDR < 0.01) and differentially methylated regions (DMR) are shown in red and blue for increased and reduced signal in PC7 relative to IMR90 ^E6E7^. Two replicates for Cut&Run and ChIP-seq, and one replicate for ONT sequencing. (**D**) Relative hTERT expression levels by RT-qPCR in PC7 (PD82-91) cells transfected with control and Rad21 siRNA pool. The means of three independent experiments with three technical replicates are shown. Two-tailed paired t test. (**E**) Hi-C sequencing analysis of B-cell Acute Lymphoblastic Leukemia (B-ALL) samples reported in (*53*). The surrounding of the TERT locus (chr5:500000-2100000, GRCh38/hg38) is shown as contact map with the loop anchor marked by an arrow. (**F**) TERT promoter DNA methylation in IMR90 ^E6E7^ and PC7. Methylation of individual CpG sites are shown as points and the average methylation of a 100 base pairs (bp) window are shown as a line. ATG and THOR are the TERT start codon and the TERT hypermethylated oncological region (chr5:1,295,206-1,295,638, GRCh38/hg38), respectively. The DMR that partially overlaps with THOR and is hypermethylated in PC7 relative to IMR90 ^E6E7^ is shown in red.

Next, we investigated how the immortal postcrisis clones activate telomerase. Using bulk whole genome sequencing (fig. S4A) we did not detect any TERT promoter mutations (*44, 45*), SVs adjacent to the TERT locus (*46–49*) or TERT amplifications (*50, 51*), all of which are mechanisms frequently found in TERT-positive cancers. To address whether an altered 3D genome organization may facilitate TERT expression as previously shown for TERT promoter mutant-positive cancer cells (*52*), we performed Hi-C analysis in the parental IMR90 ^E6E7^ cells, the mortal PC45 and the four immortal postcrisis clones. Indeed, a chromatin loop spanning the TERT locus was significantly enriched in three of the four immortal postcrisis clones (PC7, PC39, PC42) compared to the parental IMR90 ^E6E7^ but was absent in the mortal PC45 postcrisis clone (Fig. 3B). The loop comprises the T-INT1 region (chr5:1,555,972-1,558,643, GRCh38/hg38) that has been previously reported to serve as distal TERT enhancer exclusively in TERT promoter mutant-positive cells. As the isolated postcrisis clones have no TERT promoter mutations which generate novel E-Twenty-Six (ETS) transcription factor binding sites (*44, 45, 52*), we rationalized that the mechanism of TERT activation must be different. Therefore, we speculated that in the immortal postcrisis clones epigenetic changes may facilitate the loop formation, T-INT1 recruitment and TERT expression. ChIP-seq analysis in PC7 postcrisis and IMR90 ^E6E7^ cells for histone H3K27 acetylation (H3K27ac), a marker of active enhancers, showed that H3K27ac is enriched in the proximity of T-INT1 and at both loop anchors in PC7 postcrisis cells relative to IMR90 ^E6E7^ (Fig. 3C). Further, Cut&Run analysis revealed that in PC7 cells the repressive H3K27 trimethylation (H3K27me3) mark is depleted in a region upstream of the right loop anchor. Of note, this region partially overlaps with T-INT1 (Fig. 3C), suggesting that the T-INT1 enhancer may be blocked in the parental IMR90 ^E6E7^, but becomes activated in PC7 to promote TERT expression. Next, to test whether the 3D genome organization affects TERT expression, we depleted the cohesin subunit RAD21 (fig. S9B) and measured TERT expression by RT-qPCR (Fig. 3D). TERT expression levels were reduced to 50% in RAD21 depleted PC7 cells relative to control cells suggesting that the cohesin-mediated 3D genome organization partially contributes to TERT expression and consequently immortalization in the isolated postcrisis clones. Interestingly, similar loops could be detected in several published Hi-C analysis of B-cell Acute Lymphoblastic Leukemia (B-ALL) samples (*53*) (Fig. 3E). Thus, looping in the absence of TERT promoter mutations facilitates TERT expression in the immortal postcrisis clones, and in tumors such as B-ALL (Fig. 3E).

Hypermethylation of the TERT Hypermethylated Oncological Region (THOR), a part of the TERT promoter (chr5:1,295,206-1,295,638, GRCh38/hg38), is frequent in cancer and is suggested to prevent binding of negative TERT regulators (*54–56*). To address THOR methylation in IMR90 ^E6E7^ parental cells and the postcrisis clone PC7 we performed methylation-sensitive Oxford Nanopore Technologies (ONT) whole genome sequencing. Whereas methylation in PC7 was reduced genome-wide, the THOR region was hypermethylated in PC7 relative to IMR90 ^E6E7^ parental controls (Fig. 3F). Thus, THOR hypermethylation may further contribute to TERT expression and telomere maintenance in the PC7 postcrisis clone. Together, THOR hypermethylation and loop formation in the absence of TERT promoter mutations represent an additional epigenetic mechanism to upregulate TERT and acquire immortality in vitro and in vivo.

### Postcrisis clones, while escaping crisis with near diploid genomes, subsequently evolve toward tetraploidy and increased genomic instability

Chromosome counts from metaphase spreads (fig. S10A) and copy-number profiles derived from bulk whole-genome sequencing (fig. S4A) revealed subtetraploidy in late-passage cells of three out of four immortal postcrisis clones. In contrast, all mortal postcrisis clones maintained a near diploid genome (fig. S4A and S10A), opening the possibility that tetraploidization facilitates immortalization and crisis escape. To test at which ploidy these three immortal postcrisis clones escaped crisis, we performed DNA content analysis of the mortal postcrisis clones and an early and late passage of the immortal postcrisis clones (Fig. 4A and fig. S10B). Interestingly, all postcrisis clones escaped crisis with a near diploid genome (Fig. 4A and fig. S10B), indicating that TMM activation and crisis escape are not linked to tetraploidization in the isolated postcrisis clones. Instead, DNA content analysis suggested that three out of four immortal postcrisis clones acquired a subtetraploid genome during subsequent culture (Fig. 4A). These findings imply that independent whole-genome duplication events occurred in each of these three immortal clones, generating subclones with a growth advantage that ultimately dominated the respective populations.

**Fig. 4.**
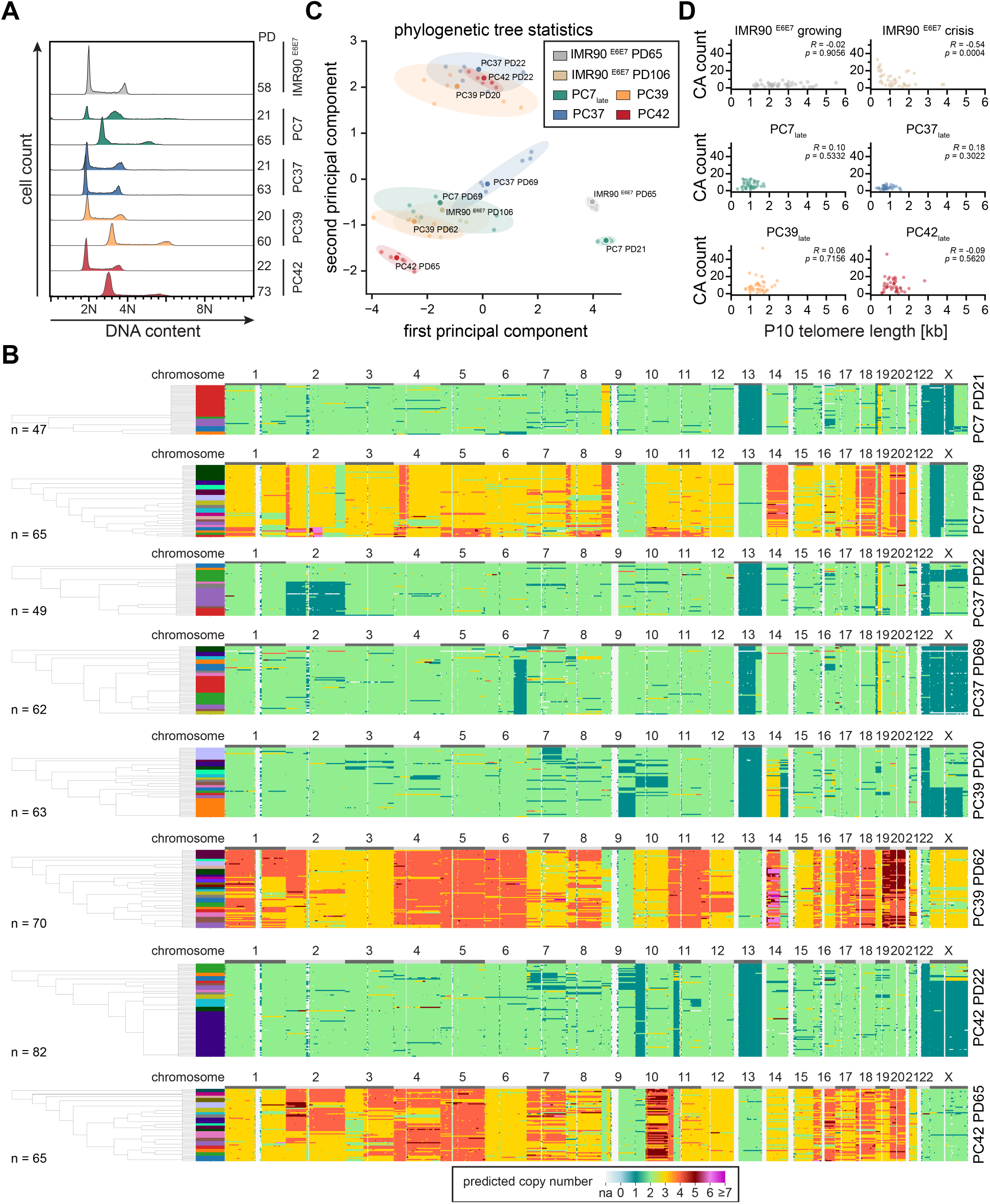
Immortal postcrisis clones evolve after crisis escape. (**A**) DNA content analysis of indicated samples. For each postcrisis clone an early and late population doubling (PD) are measured. IMR90 ^E6E7^ PD58 represent parental diploid control cells. Three independent experiments. (**B**) Copy number estimates of individual immortal postcrisis clones at an early and later PD based on single-cell Strand-seq of the indicated samples. Phylogenetic trees were calculated with PICASSO and the most representative tree of eight PICASSO runs is shown. (**C**) Principal component analysis of phylogenetic tree statistics for indicated samples. Analysis is based on eight PICASSO runs. Individual PICASSO runs are shown as points with the run closest to the center of the centroid shown as the biggest point. (**D**) Scatter plot of the recent copy number alterations against the 10^th^ percentile (P10) telomere length in kilobases (kb) for growing and crisis IMR90 ^E6E7^ cells and late passage of immortal postcrisis clones. Spearman’s rank correlation was performed.

To characterize the genetic evolution of the immortal postcrisis clones, we performed single-cell Strand-seq on an early (PD20-22) and late passage (PD62-69) of the four immortal postcrisis clones. In accordance with DNA content analysis, all single cells of the immortal postcrisis clones were near diploid at the early passages. At the late passages only PC37 maintained its near diploid ploidy state, while PC7, PC39 and PC42 cells underwent whole genome doubling (Fig. 4B). Karyotype consensus copy number data of both early and late immortal postcrisis cells included almost exclusively large-scale gain and losses, without more complex CAs such as chromothripsis or high focal amplifications at subclonal level (Fig. 4B). The most complex subclonal CAs present in the immortal postcrisis clones were detected in a subpopulation of PC7 PD69 and PC39 at chromosome 2 and 14, respectively, and these events are indicative of past BFB cycles (Fig. 4B and fig. S10C,D). As these CAs are shared by multiple individual cells in the respective postcrisis clones, we speculate that the event could be stabilized by telomerase and then propagated. At the single-cell level, whereas early postcrisis clones resembled growing pre-crisis cells, with simple karyotypes and limited heterogeneity, late postcrisis populations accumulated extensive subclonal heterogeneity despite maintaining a similarly simple consensus karyotype. This spontaneous diversification, which occurred in the absence of selective pressure and was largely undetectable by bulk DNA sequencing, prompted phylogenetic reconstruction of post crisis evolution.

To this end, we used Phylogenetic Inference of Copy number Alterations in Single-cell Sequencing data Optimization (PICASSO) (*57*) to resolve the phylogenetic trees of individual IMR90 ^E6E7^ growing, crisis and postcrisis cells based on the single-cell Strand-seq copy number predictions (Fig. 1C, 4B,C and fig. S11A-I). PICASSO analysis of IMR90 ^E6E7^ PD65 growing cells revealed one major subpopulation representing 64% of all analyzed cells (Fig. 1C). In contrast, the IMR90 ^E6E7^ crisis population structure was highly fragmented with the largest effective clone number of all samples (Fig. 1C and fig. S11G). The effective clone number was decreased in the early passages of the immortal postcrisis clones relative to IMR90 ^E6E7^ crisis cells, and phylogenetic trees were comprised of one or two major subpopulations, (Fig. 4B,C and fig. S11A-I). In the later passages of PC7, PC39 and PC42 that had undergone whole genome duplication, the effective clone number increased (fig. S11G) and their subclonal architectures became more diverse and heterogeneous, indicating a second peak in chromosome instability (fig. S11C-I). In PC37 which maintained a near diploid ploidy, the effective clone size, as well as the heterogeneity and diversity of the phylogenetic tree only increased slightly relative to the early passage, highlighting whole genome duplication as a strong accelerator of genome evolution (*2, 58, 59*). However, PC37 single-cell analysis displayed evidence of a population sweep as a subclone exhibiting a chromosome 2 disomy and chromosome X monosomy, present in 18% of PC37 PD22 cells, took over the majority of population at PD69, suggesting ongoing selection during this time of growth.

Principal component analysis of aggregated phylogenetic tree statistics revealed clustering mainly organized around karyotype heterogeneity and chromosome instability. Accordingly, highly heterogeneous and fragmented late passage postcrisis clones are grouped together and partially overlap with IMR90 ^E6E7^ crisis cells, while the early passages of PC37, PC39 and PC42 form their distinct group (Fig. 4C). Of note, the near diploid PC37 PD69 was more similar to PC37 PD22 and growing IMR90 ^E6E7^, whereas PC42, likely driven by high heterogeneity scores, was even more distinct to its early passage or growing IMR90 ^E6E7^ (Fig. 4C and fig. S11E,F,H). Importantly, while chromosome instability and heterogeneity in the late passages of the immortal postcrisis clones reach values extremely close to crisis levels (fig. S11A-I), their continued growth (Fig. 2B) suggests that chromosome instability is now tolerated by the cells. Interestingly, PC7 PD21 with its particularly simple subclonal architecture clusters together with stable, growing IMR90 ^E6E7^ (Fig. 4C). This difference may resemble the younger evolutionary age of PC7 PD21 relative to the other postcrisis clones due to the more recent clonal selection which occurred in the additional growth plateau that followed crisis (Fig. 2B).

Finally, to determine whether the reemergence of chromosome instability was primarily driven by critically short telomeres (Fig. 2C,D), we plotted the number of recent copy number alterations against the 10^th^ percentile of shortest telomeres for each chromosome arm and assessed the potential correlation using Spearman’s rank correlation coefficient (Fig. 4D, Data S1). While in IMR90 ^E6E7^ crisis cells the number of recent copy number alterations were inversely correlated with telomere length (Spearman’s R = −0.54; *p* = 0.0004), no correlation was detectable in growing IMR90 ^E6E7^ cells and the late passage of the immortal postcrisis clones (Fig. 4D), suggesting that once clones escape replicative crisis and acquire a TMM, telomeric damage is no longer the principal determinant of chromosome-specific SV formation. Therefore, after cells escaped crisis and acquired immortality, immortal postcrisis clones evolve progressively, and this evolution is accelerated by whole genome duplication events and the ability to stabilize and propagate potential detrimental SVs (fig. S11J).

## Discussion

Here, we have followed the evolution of checkpoint-defective cells through replicative crisis to spontaneously escaping postcrisis clones and immortality. Our single-cell analysis revealed high and heterogeneous genomic instability with complex CAs in crisis (Fig. 1, 4C, and fig. S2, S3, S10C,D, S11). Further, the number of terminal CAs per chromosome arm was negatively correlated with chromosome arm-specific telomere length (Fig. 1G) indicating that telomere length-dependent dysfunction is the major driver of genomic instability in crisis. In contrast, CAs were not correlated with telomere length in the immortal postcrisis clones (Fig. 4D), despite their short telomeres (Fig. 2C,D, 4D and fig. S5) and residual markers of chromosome instability (fig. S6A-C). This could be a consequence of low telomerase activity in the postcrisis clones and the preference of telomerase to elongate and stabilize those chromosome arms with the shortest telomeres (Fig. 3A, fig. S5E,F and Data S1), causing overall more homogenous telomere length distributions and lack of correlation between CA and telomere length in the immortal postcrisis clones (Fig. 4D and Data S1). Accordingly, telomere length in cancer cells is frequently shorter than in matching normal tissue (*51*), yet is sufficient to support indefinite growth. This suggests that in both the immortal postcrisis clones and cancer the TMM activity must only be sufficient to maintain the shortest telomeres, which are most prone to induce telomere-dependent growth barriers (*19, 60–62*). Additionally, a recent single-cell study of immortal micronuclei-positive MCF10A cells found elevated terminal CAs at chromosome arms with short telomere lengths, highlighting that even under conditions of an active TMM telomere dysfunction promotes spontaneous genomic instability (*35*). Thus, the short telomeres likely contribute to genome evolution in the immortal postcrisis clones and in some cancer, however, the events may be too infrequent to resolve them without phenotypic enrichment.

In contrast to crisis cells (Fig. 1C-G and fig. S2) and in accordance with a previously published whole genome sequencing analysis of eight postcrisis clones isolated in different experiments with similar setup (*63*), the emerging postcrisis clones had relatively few and simple SVs (Fig. 4B and fig. S4, S11A-I). This indicates that at least in the absence of a TMM, complex SVs are detrimental and those events are efficiently counter-selected during crisis. Consequently, cells that overcome this selective bottleneck tend to retain relatively quiet genomes. This enabled their continued proliferation and thus increased the likelihood to acquire additional alterations including TMM activation. While TMM activity grants replicative immortality, the other alterations may promote resistance to crisis cell death and support growth for additional population doublings (*17, 18*). Indeed, the growth characteristics of isolated mortal postcrisis clones and especially PC7 (Fig. 2B) suggests that growth beyond crisis can occur with no or insufficient TMM at which time a TMM can be activated (Fig. 3A and fig. S9A). However, we were unable to attribute the extended proliferation of mortal postcrisis clones to alterations in specific genes, so it is possible that their growth is driven by polygenic effects. Additionally, given the telomere length heterogeneity at every chromosome arm (Fig. 1B, 2D, and Data S1), individual cells in the population will differ in their maximal replicative age (*40, 64*), providing some cells in the population additional growth advantages. Therefore, the mortal postcrisis clones may originate from a subpopulation of IMR90 ^E6E7^ cells that had overall longer telomeres or relatively longer telomeres at the chromosome arms with the shortest telomeres (Fig. 1B). Strikingly, comprehensive TMM profiling of 976 cancer cell lines revealed that TMM was undetectable in 1.2% of the tested cancer cell lines (*65*) and the growth of some TMM-negative cancer cell lines is supported by very long telomeres (*66, 67*). Further, in neuroblastoma, a tumor found in infants and young children, which both have still relatively long telomeres (*68*), an active TMM is the highest risk factor for a detrimental prognosis, and conversely, tumors lacking a TMM have a favorable prognosis with frequent spontaneous regressions (*69*). Thus, long telomeres or genetic alterations can result in extended replicative lifespan and detectable tumors. However, as demonstrated in the mortal postcrisis clones, those tumors are ultimately restricted by telomere-dependent barriers and must sufficiently activate a TMM to achieve replicative immortality.

To maintain their telomeres, the isolated immortal postcrisis clones utilize telomerase (Fig. 3 and fig. S9A), with three immortal postcrisis clones showing an altered 3D genome organization at the TERT locus that potentially facilitates communication between the T-INT1 enhancer region and the TERT promoter. T-INT1 recruitment has been previously reported to facilitate TERT expression through transcription factor mediated looping exclusively in TERT promoter-mutant cells (*52*). However, the immortal postcrisis clones and several B-ALL samples lack TERT promoter mutations but display loop structures (Fig. 3B,E). Thus, our data suggests that independent of TERT promoter mutations, epigenetic changes can lead to THOR hypermethylation (Fig. 3F) (*54–56*) and an altered 3D organization of the TERT locus (Fig. 3B-D) that promote TERT expression and replicative immortality in the in vitro isolated immortal postcrisis clones and in patient-derived B-ALL cancer samples.

Finally, we here propose a model for crisis and crisis escape in relationship to chromosome instability during the early stages of tumorigenesis (ig. S11J): During crisis, progressive telomere length-dependent dysfunction drives extensive genomic instability, giving rise to complex CA and pronounced genomic heterogeneity. In the absence of a TMM, however, crisis cells are largely unable to stabilize chromosome ends generated by terminal breaks. These cells therefore remain susceptible to recurrent BFB cycles initiated by telomere fusions, which progressively compromise cellular fitness and are subject to strong negative selection. Telomere dysfunction thus constitutes the major selective pressure during crisis, which can be overcome through acquisition of a TMM and restoration of chromosome end protection. The rare, individual cells that gain TMM activity stabilize or even elongate their shortest telomeres. This reduces the emergences of telomere fusions with subsequent BFB cycles, facilitates crisis escape and grants replicative immortality. In addition, TMM activity can stabilize DNA double strand breaks and the outcomes of complex CAs by *de novo* telomere addition (*70–73*) allowing cell survival and proliferation in conjunction with chromosomal instability. However, as *de novo* telomere addition is dependent on the structure of the break (*70*) and TMM activity (*71, 72, 74*), cells escaping crisis with minimal or low TMM activity will by and large not have this ability. Consequently, rather than inheriting complex genomes from crisis, postcrisis clones emerge with relatively simple karyotypes and few SVs.

Post crisis, these immortal cells undergo progressive karyotype remodeling, as the counter-selective pressures that constrain genome evolution during crisis are reduced. This remodeling can be driven by multiple CA-generating processes, including mitotic errors that cause chromosome missegregation and micronucleus formation and, over successive cell divisions, promote the accumulation of additional CAs (*35, 75*). Whole-genome duplication, which has previously been linked to increased genomic instability (*2, 58, 59*), may further expand the capacity of postcrisis cells to tolerate and acquire karyotypic changes. Indeed, we observed distinct chromosome instability processes occurring post crisis, and the fixation of their respective genomic outcomes in cell subsets during the evolution from early to late passage: whole genome doubling in three out of four clones, a clonal sweep driven by whole-chromosome changes in PC37, and two BFB-like events one appearing *de novo* in PC7 and another one undergoing remodeling and divergence in PC39. During tumorigenesis, progressive chromosomal instability may enable immortal cells to acquire additional cancer hallmarks and increase their TMM activity, thereby promoting the subsequent survival and propagation of cells acquiring complex CAs. Interestingly, a recent study reported that TERT alterations predispose dysplastic nodules to malignant transition during early hepatocellular carcinogenesis (*76*), with copy-number alterations appearing only later in the course of malignant progression. Furthermore, in older individuals with shorter telomers, TERT promoter mutations are often found in dysplastic nevi, a potential precursor of melanoma (*77*). This suggests that TMM activation represents a critical early step during human tumorigenesis which not only helps to overcome replicative boundaries and achieve replicative immortality but may also facilitate the stabilization and propagation of SVs. In the human body, ongoing spontaneous genome instability in conjunction with other in vivo selection pressures, such as nutrient limitations or evasion of the immune system, may promote genome evolution with clonal selection ultimately leading to the acquisition of additional cancer hallmarks (*1*) and more complex genomes that are frequently observed in highly aggressive cancers (*7*). Taken together, our results place telomerase-mediated crisis escape as a pivotal event for karyotype evolution. This enables subsequent tolerance to CAs by stabilization of broken chromosome ends, thus permits the continued accumulation of chromosome instability and the acquisition of extensive karyotypic heterogeneity promoting malignant progression.

### Limitations of the study

Remaining limitations of our study are intrinsic to the crisis escape experiment: First, given the low rate of spontaneous crisis escape (*23*), a very large number of crisis cells is required for the experiment leading us to perform it just once. Consequently, the number of isolated clones is relatively small and no ALT-positive postcrisis clone could be isolated. Second, in the absence of TMM, replicative lifespan is restricted by telomere-dependent barriers, thus, cellular material for the mortal postcrisis clones is sparse. Finally, in our in vitro experiment, TMM acquisition overcomes the prime crisis selection pressure - telomere length-dependent dysfunction - and promotes crisis escape. However, crisis cells in the human body may simultaneously experience other selection pressures, such as immune surveillance, nutrient deprivation and hypoxia (*1*). During human tumorigenesis, these are likely to represent additional evolutionary bottlenecks not modeled in our in vitro crisis model. The high genomic instability observed in crisis cells may facilitate adaptation and resistance to these additional in vivo selection pressures and help to acquire cancer hallmarks, yet would promote clearance of crisis cells by the immune system. Finally, a recognized technical limitation of Strand-seq is its inability to reliably detect SVs smaller than 200 kb in single cells (*33*). However, relatively few established cancer-driving SVs fall within this SV size range (*78*).

## Acknowledgments

We thank members of the Karlseder lab for input. This work was supported by the Waitt Advanced Biophotonics Core Facility of the Salk Institute (RRID:SCR_014838) with funding from NIH-NCI CCSG P30 CA014195, NIH-NIA San Diego Nathan Shock Center P30 AG068635, and the Waitt Foundation, by the Flow Cytometry Core Facility of the Salk Institute (RRID:SCR_014839) with funding from NIH-NCI CCSG P30 CA014195, and Shared Instrumentation Grants S10-OD023689 (Aria Fusion cell sorter), and The Razavi Newman Integrative Genomics and Bioinformatics Core Facility of the Salk Institute (RRID:SCR_014842 and SCR_014846) with funding from NIH-NCI CCSG P30 CA014195, NIH-NIA San Diego Nathan Shock Center P30 AG068635, the NIH-NIA Liver Cancer P01 AG073084-04, the Howard and Maryam Newman Family Foundation and the Helmsley Trust.

## Funding

European Molecular Biology Organization Long Term Fellowship (ALTF668-2019) (TTS)

National Institutes of Health grant R01AG077324 (JK) Glenn Foundation for Biology of Aging Research (JK, TTS)

The American Heart Association and The Paul G. Allen Frontiers Group 19PABH134610000 (Salk Institute, JK)

National Institutes of Health grant P30CA014195 (Salk Institute NCI Cancer Center, JK) Donald and Darlene Shiley Chair (JK)

## Diversity, equity, ethics, and inclusion [optional]

We welcome statements pertaining to diversity, equity, ethics, and inclusion, as outlined in our Editorial Policies.

## Data, code, and materials availability

All data, code, and materials used in the analysis are archived at the Salk Institute and European Molecular Biology Laboratory. Sequencing data will be deposited prior acceptance.

## Supplementary Materials

### Materials and Methods

#### Cell culture

IMR90 (CCL-186) and WI38 (AG06814-N) fibroblasts were purchased from ATCC and the Coriell Institute for Medical Research, respectively. HeLa LT cells were subcloned from HeLa 1.2.11 (*79*) in the Karlseder laboratory. All cells are grown at 37 °C, 7.5% CO_2_ and 3% O_2_. Human fibroblasts were grown in GlutaMax-DMEM (Gibco, 10569-010) supplemented with 0.1 mM nonessential amino acids (Corning, 25-025-Cl), 15% fetal bovine serum (FBS) (VWR, 97068-085 Lot 323B20), 100 IU/ml Penicillin and 100 μg/ml Streptomycin (Corning, 30-002-CI). HeLa LT cells were cultured in GlutaMax-DMEM supplemented with 0.1 mM nonessential amino acids, 10% FBS, 100 IU/ml Penicillin and 100 μg/ml Streptomycin. Population doublings (PD) are calculated at each passage using the following equation: PD = ln(number of counted cells/number of initial cells)/ln(2). Cells are tested and found to be free of mycoplasma.

#### Postcrisis clone isolation experiment

IMR90 ^E6E7^ cells were grown to crisis. 81 million crisis cells were seeded at a density of 6 x 10^5^ cells per 15 cm plate (in total 135 x 15 cm plates). Media was changed every three days and plates were manually inspected for the appearance of postcrisis clones. Appearing colonies were marked and 60 days after seeding clones were isolated using clonal cylinders. Isolated clones were expanded from 96-well plates to 10 cm plate format, counted and population doubling set to zero. Growth curve analysis of postcrisis clones was performed and a frozen stock was generated regularly.

#### siRNA transfection

siRNA transfections were performed using Lipofectamine RNAiMAX kit (Thermo Fisher Scientific, 13778030). Cells were transfected with 30 nM siRNA on two consecutive days and cells harvested 72 h after first transfection. Dharmacon ON-TARGETplus SMART siRNA pools (Horizon) and have the 5’-3’ target sequences: Non-targeting pool (NT) (D-001810-10): UGG UUU ACA UGU CGA CUA A, UGG UUU ACA UGU UGU GUG A, UGG UUU ACA UGU UUU CUG A, UGG UUU ACA UGU UUU CCU A; RAD21 (L-006832-00): GCU CAG CCU UUG UGG AAU A, GGG AGU AGU UCG AAU CUA U, GAC CAA GGU UCC AUA UUA U, GCA UUG GAG CCU AUU GAU A.

#### Retroviral and lentiviral transduction

Retro- and Lentivirus were produced in the laboratory. To produce retroviruses in Phoenix cells, calcium phosphate transfection with 100 μM chloroquine was used as described previously (*17*). To produce lentivirus, Lenti-X 293T cells (Takara, 632180) were transfected with 7 μg plasmid DNA using Lenti-X Packaging Single-Shot system (Takara, 631276). Media was changed after 24 h and virus containing supernatant collected 48 and 72 h after transfection. Supernatant was concentrated using Lenti-X concentrator (Takara, 631232) following the manufacturer’s protocol. Lentivirus was resuspended in fresh media and added to the cells in the presence of Lenti-Blast (OzBiosciences, LB00500).

#### Soft-agar assay

Soft agar assay was performed in 6 well plates as described in (*80*). 5000 cells per well were imbedded in growth media with 0.3% agar and layered on top of growth media with 0.5% agar. Cells were grown for 21 days, stained with nitro blue tetrazolium chloride solution (Invitrogen, N6495) and imaged with G:Box Chemi XT4 imager (Syngene). Three independent experiments with three technical replicates were performed.

#### Western blotting

Western blots were performed as described previously (*81*). In brief, cells were washed with 1x DPBS and lysed in NuPage LDS sample buffer (Invitrogen, NP0007) containing 2% (v/v) benzonase (Roche) at 1 x 10^4^ cells per μL. Proteins were denatured at 95 °C for 5 min, separated on NuPAGE Bis-Tris gels (Invitrogen, NP0321, NP0322 and NP0336) in either NuPAGE MES or MOPS SDS running buffer (Invitrogen, NP0002 and NP0001, respectively) and transferred to a nitrocellulose membrane (Amersham, 10600037) at 110 V at 4 °C for 1h. Membranes were blocked with either 5% milk/TBST or 5% BSA/TBST at RT for 1 h and incubated with primary antibody (table S2) at 4 °C, overnight. Horseradish peroxidase (HRP) linked anti-mouse IgG (CST, 7076) and anti-rabbit IgG (CST, 7074) were used as secondary antibodies. Blots were developed using ECL kits (Prometheus, 20-301B and 20-302B) and imaged with the G:Box Chemi XT4 imager (Syngene).

#### Immunofluorescence

Cells were seeded on glass cover slips 24 h prior the experiment. Cells were fixed in 4% paraformaldehyde (Electron Microscopy Sciences, 15710) in 1x PBS for 10 min, washed with PBS three times, permeabilized in 0.3% Triton-X100/PBS at RT for 10 min and incubated in blocking solution (5% BSA in PBS) at RT for 1 h. Samples were then incubated with primary antibody either at RT for 1 h or 4 C overnight. After three PBS washes, samples were incubated with AlexaFluor 488 goat anti-rabbit IgG (H+L) (Thermo Fisher Scientific, A-11034) and AlexaFluor 568 goat anti-Mouse IgG (H+L) (Thermo Fisher Scientific, A-11004). Finally, samples were washed with PBS three times and mounted in ProLong Diamond with DAPI (Invitrogen, P36971). Samples were imaged using either the Zeiss LSM 880 with the Airyscan or the Zeiss Axio Imager.M2 epifluorescence microscope. ZEN (Zeiss) and Fiji (ImageJ) were used for image analysis.

#### Metaphase spreads for quantification of chromosome fusions

Metaphase spreads were prepared as described in (*82*). In brief, cells were enriched in metaphase with 200 ng/ml colcemid (Thermo Fisher Scientific, 15212012 Gibco KaryoMAX) for 3 h, collected by trypsinization and centrifuged at 270 x g for 4 min. Cells were swollen in hypotonic solution (75 mM KCl) at 37 °C for 7 min. To prefix cells, 1 mL of freshly prepared fixative solution (3:1 (v/v) methanol: glacial acetic acid) was added to samples, mixed by inversion and samples pelleted at 270 x g for 4 min. Cell pellets were resuspended by tapping and 10 ml fixative solution added to samples in a dropwise fashion. Next, cells were washed twice with 10 ml fixative solution before samples were dropped on superfrost plus slides (VWR, 48311-703). Slides were dried overnight and rehydrated in PBS for 5 min and fixed in 3.7% formaldehyde (Fisher scientific, BP531). Samples were washed three times with PBS and then dehydrated by an ethanol bath series (70%, 90% and 100% ethanol, 3 min each) and air-dried. Slides were layered with 50 μl of 0.3 ng/μl TelC-Alexa488 PNA probe (PNA Bio, F1004) and 0.3 ng/μl CENT-Cy3 PNA probe (PNA Bio, F3003) diluted in hybridization buffer (70% (v/v) deionized formamide; 0.25% (v/v) blocking reagent (Roche, 11096176001, dissolved in 100 mM malic acid, 150 mM NaCl, pH 7.5); 10 mM Tris pH 7.5, 5 mM MgCl_2_), covered with cover glass, denatured at 80 °C for 5 min and incubated at RT in the humidity chamber for 2 h. Next, while shaking, slides were washed twice with PNA wash A (70% (v/v) formamide,10 mM Tris-HCl, pH 7.5) for 5 min each, followed by three washes of PNA Wash B (150 mM NaCl, 50 mM Tris-HCl, pH 7.5, 0.08% Tween-20) for 5 min each, with 100 ng/ml DAPI (Sigma, D9542) in the second wash. Slides were dehydrated and air-dried as described above and mounted in ProLong Diamond (Invitrogen, P36970). Metaphases were imaged using either Metafer or Zeiss Axio Imager.M2 microscope with 63x oil objectives.

#### Detection of Telomere Dysfunction-Induced Foci on Metaphase Spreads (MetaTIFs)

The MetaTIF assay was performed as previously described (*83*). Briefly, cells were seeded on 10-cm dishes. To enrich for mitotic cells, the cultures were treated with 20 ng/ml Colcemid (Thermo Fisher Scientific, 15212012 Gibco KaryoMAX) for 1 hour. Cells were harvested by trypsinization, resuspended in a hypotonic buffer solution (0.2% (w/v) KCl and 0.2% (w/v) sodium tricitrate) at a density of 1 x 10^5^ cells/ml, and incubated for 7 minutes at room temperature (RT). Metaphase spreads were prepared by cytocentrifugation (Cytospin 3, Shandon) onto Superfrost Plus slides (VWR, 48311-703) at 2,000 rpm for 10 minutes with medium acceleration. Slides were fixed in 4% (v/v) formaldehyde in PBS for 10 minutes at RT, followed by blocking in Immunofluorescence Blocking Buffer (Cell Signaling Technology, #12411) supplemented with 100 μg/ml RNase A (Sigma) for 15 minutes at 37 °C. For IF, slides were incubated overnight at 4 °C with a primary antibody against γH2AX (Millipore, clone JBW301) diluted 1:1,000 in Immunofluorescence Antibody Dilution Buffer (Cell Signaling Technology, #12378). After washing in PBS, slides were incubated for 1 hour at RT with an Alexa Fluor 594-conjugated anti-mouse IgG (H+L) F(ab’)2 fragment secondary antibody (1:1,000; Cell Signaling Technology, # 8890). Following IF, slides were fixed again in 4% (v/v) formaldehyde in PBS for 10 minutes at RT, rinsed in Milli-Q water, and dehydrated through a graded ethanol series (70%, 90%, and 100% for 2 minutes each). Telomeric DNA was labeled via fluorescence in situ hybridization (FISH) using 0.3 ug/ml Alexa 488-OO-(CCCTAA)_3_ PNA probe (PNA Bio, F1004) in hybridization buffer (70% (v/v) deionized formamide, 10% Roche blocking reagent, and 10 mM Tris-HCl, pH 7.5). Samples were denatured at 80 °C for 5 minutes and hybridized overnight at RT. Slides were washed twice in PNA Wash A (70% (v/v) formamide, 10 mM Tris-HCl, pH 7.5) and three times in PNA Wash B (50 mM Tris-HCl, pH 7.5, 150 mM NaCl, and 0.08% (v/v) Tween-20). 50 ng/ml DAPI (Sigma, D9542) was included in the second Wash B for nuclear counterstaining. Slides were rinsed in Milli-Q water, dehydrated in ethanol, air-dried, and mounted with ProLong Diamond Antifade Mountant (Invitrogen, P36970).

Images were acquired using a Zeiss Axio Imager.M2 microscope equipped with a 63x/1.4 NA oil immersion DIC objective. For each metaphase spread, five Z-stacks were acquired and processed as maximum intensity projections. MetaTIFs, defined as γH2AX signals colocalizing with PNA-labeled telomeres, were manually scored using Fiji (ImageJ). A total of 16 metaphases (n = 8 per replicate) were analyzed per sample.

#### Fusion analysis

Single-molecule telomere fusion PCR assays were performed as previously described (*19, 84*). 50 ng of phenol/chloroform extracted DNA were subjected to PCR using the following primers: 21q1 (CTTGGTGTCGAGAGAGGTAG), 17p6 (GGCTGAACTATAGCCTCTGC) and XpYpM (ACCAGGTTTTCCAGTGTGTT). Six reactions per sample were set up each containing 0.5 μM PCR primers and 0.5 U of a 10:1 mixture of Taq (Thermo Fisher Scientic) and Pwo polymerase (Roche). The fusion amplicons were resolved by TAE agarose gel (0.5%) electrophoresis and detected by Southern hybridization with random-primed [α-33P] labelled (Hartmann Analytic GmbH) telomere-adjacent probes together with probes to detect 1 kb (Stratagene) and 2.5 kb (BioRad) molecular weight markers, followed by phosphorimaging with a Typhoon FLA 9500 (Cytiva).

#### STELA

STELA analysis was performed as described previously (*19, 40*). Briefly, 10 ngs of phenol/chloroform extracted DNA was added to 40 μL 10 mM Tris–HCl (pH 8) and Telorette2 linker (0.25 μM). Four 10 μL PCR reactions were set up separately for the XpYp and 17p telomeres each containing 1 μL of DNA/Telorette2 solution, 0.5 μM of telomere-specific primers (17pseq1rev: 5′-GAATCCACGGATTGCTTTGTGTAC-3′) or (XpYpE2: 5′-TTGTCTCAGGGTCCTAGTG-3′), 0.5 μM Teltail primer and 0.5 U of a 10:1 mixture of Taq (Thermo Fisher Scientific) and Pwo polymerase (Roche). PCR amplicons were resolved by 0.5% TAE agarose gel electrophoresis and detected by Southern hybridization with a random-primed [α-33P] labelled (Hartmann Analytic GmbH) TTAGGG repeat probe together with probes to detect 1 kb (Stratagene) and 2.5 kb (BioRad) molecular weight markers, followed by phosphorimaging with a Typhoon FLA 9500 (Cytiva) and analyzed using ImageQuant TL (Cytiva).

#### Telomere sequencing (Telo-seq)

Telo-seq was performed as previously described in (*32*) with modifications for the R10 flow cell chemistry and multiplexing according to Oxford Nanopore Technologies (ONT)’ instructions. In brief, high-molecular weight genomic DNA (gDNA) was extracted using Monarch HMW DNA extraction Kit for Cells and Blood (NEB, T3050) according to manufacturer’s instructions and quantified using the Qubit dsDNA BR assay (Invitrogen, Q32853).

Telo-adapter oligonucleotides (TA1-6, 40 μM final concentration, table S3) were individually hybridized with the corresponding Telo-splint oligonucleotides (TS1-6, 40 μM final concentration, table S3) in annealing buffer (50 mM NaCl, 10 mM Tris HCl, pH 7.5, 1 mM EDTA final concentration) using a thermocycler (BioRad, C1000) (5 min 95 °C denaturation followed by gradual −0.1 °C/cycle decrease in temperature for 850 cycles). Annealed adapters were diluted to a final concentration of 1.33 μM in TE buffer (10 mM Tris-HCl, pH 8.0, 0.1 mM EDTA). For each sample, in a final volume of 200 μl, 15 μg gDNA was mixed with 1 mM ATP (Thermo Scientific, R0441), the Telo-adapter (final concentration 33.25 nM), 10 U/μl Salt-T4 DNA ligase (NEB, M0467) in 1x rCutsmart buffer (NEB, B6004) and incubated at 35 °C for 16 h followed by heat inactivation at 65 °C for 10 min. The reaction mixes were digested with 80 U EcoRV-HF (NEB, R3195) at 37 °C for 30 min and heat inactivated at 65 °C for 20 min. Next, the digested samples were treated with 9 μl Klenow fragment (3’-5’ exo-) (NEB, E6053) in 250 μl 1x NEBNext dA-tailing reaction buffer (NEB, E6053) at 37 °C for 30 min to perform dA tailing. 250 μl AMPure XP beads (1x v/v) (Beckman Coulter, Inc. #A63881) were added to the dA-tailing reaction, incubated at RT for 5 min, spun down on a magnet, washed twice with Long Fragment Buffer (LFB) (ONT, EXP-LFB001) and resuspended in 55 μl water. The samples were incubated at 37 °C for 15 min and spun down on a magnet. All barcoded and telo-adapted samples were combined in one tube and incubated with 100 nM Telo-splint mix (TS1-6, stock concentration 2 μM each)) in the presence of 50 mM NaCl at 50 °C for 1 h. Annealed DNA was purified using 180 μl AMPure XP beads (0.5x v/v) as described above using 66 μl water to elute DNA.

Next, 5 μl of Native Adapter (ONT, EXP-NBA114) was ligated to 65 μl of the sample using 10 μl Salt-T4 DNA ligase in 1x NEBNext Quick ligation reaction buffer (NEB, B6058) at RT for 20 min followed by purification with 50 μl AMPure XP beads (0.5x v/v), as described above using 33 μl elution buffer (ONT) to elute the library. 32 μL library was further processed according to manufacturer’s instructions and loaded on R10.4.1 PromethION flow cell (ONT, FLO-PRO114M). Library was sequenced using a PromethION 2 Solo sequencer (ONT). Six samples were multiplexed in one flow cell. R10 reads were basecalled from their respective POD5 files using dorado v1.3.0 and the dna_r10.4.1_e8.2_400bps_sup@v5.2.0 model. Telomeric reads were identified in the resulting FASTQ files using TARPON pipeline (version 2.1.1) (*85*). The TARPON-identified telomeric reads were mapped to the assembled IMR90 ^E6E7^ contigs and reads with mapping quality of 40 or higher were considered for downstream analysis. The TARPON-quantified VRR-telomere length was used as telomere length throughout this manuscript.

Data was further analyzed and plotted using R (version 4.3.1) with packages dplyr (version 1.1.4), ggplot2 (version 3.5.1), and tidyr (version 1.3.0) in RStudio (Posit Software, PBC, version 2023.12.0+369).

#### Telomeric Repeat Amplification Protocol

Telomerase activity was analyzed by telomeric repeat amplification protocol (TRAP) as previously described (*43*). In brief, cells were washed twice with ice cold PBS and collected in ice cold PBS by scraping and centrifugation (300 x g, 4 °C, 8 min). Cell pellets were resuspended in 2x telomerase buffer (TB) (20 mM Tris, pH 8.5, 6 mM KCl, 2 mM MgCl_2_, 2 mM DTT with protease inhibitor cocktail (Sigma, P8340), 100 μM PMSF (Roche, PMSF-RO), 10 μM Peptstatin A (Sigma, P5318) and 100 U RNasin (Promega, N251B)) and centrifuged (376 x g, 4 °C, 5 min). Washed cells were carefully resuspended in 1x TB (10 mM Tris, pH 8.5, 3 mM KCl, 1 mM MgCl_2_, 1 mM DTT with protease inhibitor cocktail (Sigma, P8340), 100 μM PMSF (Roche, PMSF-RO), 10 μM Peptstatin A (Sigma, P5318) and 100 U RNasin (Promega, N251B)) and incubated on ice for 15 to 20 min. Samples were homogenized by six strokes of a 25 gauge needle and a 1 cc syringe. Once cell membranes rupture, samples were incubated on ice for an additional 15 min. Extracts were centrifuged (21130 x g, 4 °C, 8 min) and supernatant was transferred to new 1.5 ml reaction tube. The glycerol concentration was adjusted to 20 % and NaCl was added to a final concentration of 0.1 M. Aliquots were flash frozen and stored at −80 °C.

Protein concentration of extracts was quantified using the Qubit Protein assay (Invitrogen, Q33211). Telomerase elongation reaction was performed with 2 μg extract in the presence of 10 μg BSA (NEB, B9001) per reaction in 1x TB containing 0.1 mM each of dATP, dGTP, dTTP (NEB, N0446), 250 μM Spermidine (Sigma, S2501), 25 μM Spermine (Calbiochem, 5677), 5 mM MgCl_2_ and 17.5 ng/μL primer #6899 (5’-GCA CAT GCA TCG AGA GTT-3’, ValueGene) at 30 °C for 30 minutes. To stop the reaction, 130 μL stop mix containing 1 mg/ml proteinase K (Millipore, 70663-4), 0.5% SDS (Sigma, 05030), 10 mM Tris, 1mM CDTA (Sigma, 319945) was added to each reaction and incubated at 37 °C for 1 h. Next, 7 μL 5 M LiCl followed by phenol-chloroform extraction and ethanol precipitation. To PCR amplify the telomerase elongation product, 2.5 μL of the telomerase elongation product was mixed on ice with PCR reaction mix (1x AmpliTaq buffer, 1.5 mM MgCl_2_, 0.2 mM of dATP, dGTP and dTTP, 2.5 μM dCTP, 0.125 μL 32P dCTP (Amersham, #AA0005), 0.55 μL Morin primer (5’-GCA CAT GCA TCG AGA GTT-3’), 0.55 μL secondary primer #6900 (5’-CGA CTT GCC TAA CCC TAA-3’) and 1 U AmpliTaq (Applied Biosystems, N8080153)) in a total volume of 25 μL. PCR was run in a thermal cycler as follows: 94 °C for 4 min followed by 29 cycles of 94 °C for 30s, 55 °C for 1 min, 72 °C for 1 min, and a final incubation at 72 °C for 5 min. The amplified products were analyzed on 8% acrylamide-urea denaturing gels. The gel was dried at 75 °C for 1 h, exposed to a PhosphoImager screen and screen scanned in a Typhoon 9400 PhosphoImager (Amersham, GE Healthcare).

#### DNA content analysis

Cells were harvested at 70% confluency by trypsinization and resuspended in 500 μL 1x DPBS (Corning, 20031-CV). While vortexing samples at medium speed, 1.5 ml ice cold absolute ethanol was slowly added to cells dropwise. Samples were stored at 4 °C. Samples were spun down (270 x g, RT, 4 min), washed once with 500 μL 1% FBS/1x DPBS and resuspended in 500 μL staining buffer (2 mM EDTA, 0.2 mg/ml RNaseA (Invitrogen, 12091039), 4 μg/ml propidium iodide (PI)(Sigma, P4864) in 1x DPBS)). Samples were incubated in the dark at 37 °C for 30 min, filtered through a cell strainer and measured using a CantoII flow cytometer (BD Bioscience)). At least 10,000 cells were recorded per sample. Analysis was done in FlowJo (BD Bioscience, version 10).

#### RT-qPCRs

Total RNA was isolated with RNeasy Mini Kit (Qiagen, 79106) with QIAshredder (Qiagen, 79656) and DNaseI (Qiagen, 74256) digestion following manufacturer’s protocol and quantified using DS11-FX Spectrophotometer / Fluorometer (DeNovix). cDNA was synthesized using SuperScript™ III First-Strand Synthesis System (Invitrogen, 18080051) following manufacturer’s recommendations with 1:1 mix of oligo-dT and random hexamer primers. For RT-qPCRs, 2 μL diluted cDNA was mixed with 0.5 μM primer mix in Power SYBR Green PCR (Applied Biosystems, A25742) in a total volume of 10 μL. Reactions were run in 384-well plates (Applied Biosystems, 4309849) in a CFX384 Touch Real-Time PCR Detection System (Bio-Rad). Relative gene expression was normalized using reference gene and calculated using the Comparative CT Method (ΔΔCT method). Primers are listed in table S3.

#### Bulk RNA sequencing

Samples were processed in parallel to the samples previously reported (*17*). Total RNA was isolated using TRIzol (Invitrogen, 15596018) and purified using the RNAeasy Mini Kit with a DNase I double digestion according to manufacturer’s instructions. The quality of the isolated total RNA was evaluated using the Agilent TapeStation 4200 and RNA-seq lirbaries were prepared with 500 ng total RNA using TruSeq stranded mRNA sample preparion kit according to the manufacturer’s protocol (Illumina). RNA-seq libraries were multiplexed, normalized and pooled for sequencing on the HiSeq400 (Illumina) as 75 bp single reads. Image analysis and base calling was performed using CASAVA v.1.8.2 (Illumina). Reads quality was tested using FASTQC (v.0.11.8) (https://www.bioinformatics.babraham.ac.uk/projects/fastqc/) and reads were mapped to hg19 human genome using the STAR aligner (v.2.5.3a) (*86*) with default parameters. Raw or transcripts per kilobase million (TPM) gene expression was quantified across all the exons of RefSeq genes with analyzeRepeats.pl in HOMER (v.4.11.1) (*87*), which used the top-expressed isoforms as a proxy for gene expression. Differential gene expression was performed on the raw gene counts with the R package, DESeq2 (v.1.24.0) (*88*), using replicates to compute within-group dispersion. Differentially expressed genes were defined as having a False Discovery Rate < 0.05 and a log2[fold change] > 0.585 (∼1.5 fold) when comparing two experimental conditions. GO enrichment analysis was performed using WEB-based Gene Set Analysis Toolkit (WebGestalt) (*89*) and the R package clusterProfiler (*90*). Data was plotted using the R packages ggplot2, ComplexHeatmap (*91*) and VennDiagram (*92*) as well as https://bioinformatics.psb.ugent.be/webtools/Venn/.

#### Bulk whole genome next-generation sequencing

For bulk whole genome sequencing by next-generation sequencing, genomic DNA was isolated with Qiagen DNAeasy Blood and Tissue Kit (Qiagen, 69504) according to manufacturer’s protocol for purification of total DNA from animal cells with spin-columns. 150 bp paired-end next-generation sequencing was performed on Illumina NovaSeq sequencer at Novogene. After removing low quality and adapter containing reads, reads were aligned to hg38 with bwa mem (v.0.7.12-r1039), and duplicate-marked with Picard (v.1.130). SNPs and INDELs were called with GATK (v.4.0.8.1)’s haplotypecaller following BQSR. Variants were hard filtered with VCFtools (v.0.1.13) to the following thresholds: max-missing = 0.4, mac = 2, min-meanDP = 15, minQ = 30. Variants deviating from the parental IMR90 ^E6E7^ PD66 genotype were annotated to genomic features using the Ensembl annotation of GRCh38v.39 (Ensembl 105). To call SVs, Manta (v.1.6.0) and Delly (v.1.0.3) were used. SVs called by both softwares were extracted and verified by hand in IGV where SVs visible in multiple distinct read fragments were retained. Copy number variants were called using ascatNGS from cgpwgs (v.2.1.0). Circos plots were generated using circlize R package (v.0.4.17) (*93*). Mutation signatures in the postcrisis clones were analyzed using SigProfilerAssignmentR with COSMIC mutational signatures (v.3.4) (*94*).

#### Bulk whole genome long-read sequencing

For ONT long-read whole genome sequencing, high-molecular weight genomic DNA was extracted using Monarch HMW DNA extraction Kit for Cells and Blood (NEB, T3050) according to manufacturer’s instructions and quantified using the Qubit dsDNA BR assay (Invitrogen, Q32853). Libraries of size-selected and Megaruptor 3 (Diagenode) fragmented genomic DNA of IMR90 ^E6E7^ PD70 and PC7 PD73 were prepared using Ligation Sequencing Kit V14 (ONT, SQK-LSK114) according to ONT’s protocol (GDH_9174_v114_revQ_10Nov2022) and sequenced on a PromethION 2 Solo sequencer (ONT). One PromethION flow cell (ONT, R10.4.1) per sample was used. Reads were basecalled in super-accuracy mode using Dorado (model dna_r10.4.1_e8.2_400bps_sup@v4.2.0). Basecalled reads were aligned to the hg38 reference genome (GRCh38 initial release, UCSC, January 2014), somatic Single Nucleotide Variant (SNV) calling and differential methylation analysis between PC7 (tumor) and IMR90 ^E6E7^ (normal) were performed using EPI2ME wf-somatic-variation v1.4.1 (commit f53d3da) with ClairS for somatic variant calling and modkit for base modification analysis. The germline VCF from the IMR90 ^E6E7^ normal sample was provided for somatic filtering. All pipelines were executed in Singularity containers using Nextflow v24.04.4.

#### Single-cell Strand-seq

We performed single-cell genomic sequencing with Strand-seq: samples were processed at large-scale using a liquid handling robotic platform with an adaptation of the one-pot StrandSeq library construction method previously described (*34, 35, 95*). Libraries were sequenced on a NextSeq5000 (MID-mode, 75 bp paired-end) followed by demultiplexing. Reads were aligned to GRCh38 reference assembly with BWA-MEM v.0.7.17, yielding a median of ∼254,000 mapped unique fragments per cell. We then processed read count data with strandtools to generate copy-number profiles and detect chromosomal abnormalities (*35*).

#### Inference of subclonal architecture

Copy-number profiles were generated from Strand-seq data using *strandtools* predictions, which were converted into a genome-wide copy-number matrix. Genomic bins with more than 20% missing predictions across cells were excluded from further analysis, as were bins overlapping centromeric regions.

For each genomic bin, the modal copy-number state across all cells was determined and used as the reference state. Deviations from this modal copy number were calculated for every cell and contiguous bins with identical deviation values were segmented. To reduce noise arising from isolated prediction errors, segments shorter than 10 Mb were removed by resetting their deviation values to zero, thereby restoring the modal copy-number state.

To identify recurrent copy-number alterations relevant for subclonal architecture, pairwise intersection-over-union (IoU) scores were calculated between all segments sharing the same deviation value. Recurrent segments were retained only if they exceeded a minimum equivalent support of two cells. The resulting filtered deviation matrix was then restricted to genomic bins containing at least one retained copy-number deviation and used as an input for phylogenetic reconstruction with PICASSO (*57*). Phylogenies were inferred using a minimum clone size of two cells, an assignment confidence threshold of 0.8, and an assignment confidence proportion of 0.9. PICASSO was executed for eight independent iterations.

For each inferred phylogeny, ancestral copy-number states were reconstructed by traversing the tree from the leaves toward the root. At each genomic bin, if one child node exhibited the modal state while the other carried a copy-number change, the parent was assigned the modal state, consistent with the assumption that the alteration occurred in the descendant lineage. If the two child nodes carried opposing copy-number changes, the parental state was considered unresolved. After the initial traversal, unresolved bins were resolved by propagating information from ancestral nodes wherever possible. Once ancestral copy-number states had been assigned throughout the tree, complete karyotypes for all internal nodes were reconstructed by combining the inferred deviations with the genome-wide modal copy-number profile. Copy-number alterations were subsequently identified by comparing each node with its immediate parent. Adjacent altered segments separated by gaps smaller than 5Mbp were merged into single events.

Finally, copy-number alterations were classified according to their phylogenetic origin. Recent alterations were defined as segments differing between individual cells (leaf nodes) and their immediate ancestral node, representing changes acquired within the current subclone. Ancestral alterations were defined as segments differing between each internal ancestral node and its parent, corresponding to copy-number events acquired earlier during clonal evolution.

#### Clonal tree analysis

To quantitatively characterize clonal architecture, we computed a set of established phylogenetic tree balance indices together with diversity and copy-number heterogeneity measures. These metrics provide complementary information on clonal architecture, evolutionary branching, and genomic diversity.

Tree balance was assessed using the Sackin and corrected Colless indices. The Sackin index was calculated as the sum of the depths of all terminal nodes (leaves), where leaf depth corresponds to the number of ancestral internal nodes separating a leaf from the root. Because the absolute Sackin index depends on tree size, values were normalized by the expected value for a perfectly symmetric tree, *n × log₂(n)*, where *n* is the number of leaves. The Colless index was computed by summing, across all internal bifurcating nodes, the absolute difference in the number of descendant leaves between the left and right subtrees. Colless values were normalized by the maximum possible imbalance for a tree of the same size, *((n − 1) × (n − 2)) / 2*, resulting in a size-independent measure of tree asymmetry.

Clonal diversity was quantified using the Shannon and Simpson diversity indices based on inferred clone sizes. The Shannon index measures the uncertainty of predicting the clonal identity of a randomly selected cell and captures both clone richness and evenness. The Simpson diversity index measures the probability that two randomly selected cells belong to the same clone and is weighted more strongly toward dominant clones.

To estimate genomic heterogeneity within individual clones, we calculated the intra-clonal heterogeneity as the mean Manhattan (L1) distance between each single-cell copy-number profile and the consensus copy-number profile of its assigned clone, by summing the absolute copy-number differences across all genomic bins and averaged across all cells belonging to the same clone.

To obtain an intuitive estimate of clonal complexity, we calculated the effective clone number, defined as the exponential of the Shannon entropy of the clone size distribution. This quantity corresponds to the number of equally sized clones that would produce the observed level of diversity and is an indication of clone fragmentation. To summarize overall evolutionary heterogeneity and tree topology, the metrics described above were scaled and summed into a composite heterogeneity score.

To further partition genomic variability across the phylogeny, copy-number differences were quantified using Manhattan distances. Total copy-number variability was defined as the mean distance between leaves and the root node. Within-clone variability was calculated as the clone size-weighted average distance between individual cells and their corresponding clone ancestor. Between-clone variability was calculated as the clone size-weighted distance between each clone ancestor and the root. The relative contributions of within- and between-clone variability were obtained by normalizing both quantities to the total variability, allowing direct comparison of genomic diversification occurring within-versus between-clone variability.

The evolutionary rate was estimated as the average amount of copy-number change accumulated along each lineage. Specifically, the Manhattan distance between every leaf and the root node was calculated, summed across all leaves, and divided by the total number of leaves in the tree.

Finally, the multidimensional tree features were summarized using principal component analysis (PCA) performed on the following variables: normalized Colless index, normalized Sackin index, Shannon diversity, Simpson diversity, intra-clonal heterogeneity, evolutionary rate, within-clone variability fraction, between-clone variability fraction, within-to-total variability ratio, and effective clone number. Because phylogenetic inference was repeated eight times for each sample, a single representative tree was selected by identifying the iteration whose feature vector was closest to the centroid of all iterations for that sample.

#### In situ Hi-C

Hi-C was performed using the Arima-Hi-C+ Kit (Arima Genomics) according to the manufacturer’s instructions. Briefly, cells were fixed for 10 minutes at room temperature in PBS and 2% methanol-stabilized formaldehyde (Fisher BioReagents, BP531), before addition of Stop Solution 1 to quench the formaldehyde. Cells were then washed twice with PBS and pellets flash-frozen for storage at −80 °C. Cells were lysed and nuclei permeabilized, followed by restriction digest and ligation with biotin fill-in. Samples were decrosslinked, digested with proteinase K, and DNA was purified using AMPure XP beads (Beckman Coulter, A63881). DNA was sonicated to ∼400 bp on a Covaris M220 (50-W power, 10% duty factor and 200 cycles per burst for 78 s) before size-selection using AMPure XP beads. Biotinylated DNA was bound to Arima Enrichment Beads and KAPA HyperPrep (Roche, KK8502) was performed on the bead-bound DNA using TruSeq DNA Single Indexes (Illumina, 20015960), according to instructions from Arima Genomics. Libraries were quantified by qPCR using KAPA Illumina Library Quantification Kit (Roche, KK4854) and Agilent TapeStation was used to estimate individual library sizes. Libraries first underwent a quality-control paired-end 37-bp run on Illumina MiniSeq, before paired-end 50 bp sequencing on Illumina NovaSeqX.

#### Hi-C Analysis

##### Alignment and processing

Hi-C data was aligned using a previously published pipeline (*96*). Briefly, read pairs were independently aligned using BurrowWheeler Aligner (BWA)-MEM to the hg38 reference genome assembly (97) before being combined post-alignment. Optical and PCR duplicates were then removed using Picard MarkDuplicates (https://broadinstitute.github.io/picard/ (*98*)). Contact matrices were generated and used to create balanced multi-resolution cooler files (.mcool) (*99*). For visualization, contact matrices from experimental replicates were merged prior to generation of the mcool file.

##### Obtaining counts and differential analysis

Read counts were extracted from the non-merged mcool files for pairwise contacts genome-wide at 25 kb resolution. Contacts between adjacent bins and bins located more than 5 Mb apart were excluded, as were contacts where the total read counts was fewer than 10. Differential analysis was then performed using the remaining contacts using edgeR (*100*).

#### IMR90 ^E6E7^ custom reference genome

After ONT whole genome sequencing of IMR90 ^E6E7^ at PD70, dorado v1.3.0 with the dna_r10.4.1_e8.2_400bps_sup@v5.2.0 model was used to basecall the resulting POD5 files. Both the ONT reads output from dorado and Hi-C reads were then incorporated into an assembly using hifiasm v0.25.0-r726. Hifiasm was run with the parameters --ont --telo-m CCCTAA --hom-cov 28, to account for the usage of ONT reads, address telomere sequences, and set the proper homozygous read coverage, respectively. The assembled contigs were mapped to the GCF_009914755.1_T2T-CHM13v2.0 human reference using minimap2 v2.30 to identify the telomeric contigs. The TARPON telomeric reads were also mapped to the assembled contigs with minimap2 to confirm the presence of matching telomeric repeat sequences. The acrocentric chromosome arms 13p,14p,15p, 21p and 22p (*101*) and chromosome arm 20p of IMR90 ^E6E7^ could not be assembled.

#### CUT&RUN

CUT&RUN was performed using the CUTANA CUT&RUN Kit (EpiCypher, 14-1048) according to manufacturer’s instructions. Briefly, 500,000 unfixed cells were bound to activated Concanavalin A beads before overnight incubation with anti-IgG (EpiCypher, 13-0042k), anti-H3K4me3 (EpiCypher cat. no. 13-0041k), or anti-H3K27me3 (Cell Signaling, 9733). The following day, cells were incubated with pAG-MNase to release bound DNA regions and *E. coli* DNA was spiked-in. Released DNA was subject to clean-up before library preparation using the CUTANA CUT&RUN Library Prep Kit (EpiCypher, 14-1001). Libraries were sequenced paired-end 75 bp on Illumina NovaSeqX or Element AVITI.

#### ChIP-seq

Approximately 10 million cells were fixed in PBS and a final concentration of 1% methanol-free formaldehyde (Thermo Scientific Pierce, 28908) for 10 minutes at room temperature, before being quenched with 125 mM glycine for 10 minutes. Cells were washed twice in PBS before harvesting, then flash frozen and stored at −80°C. Pellets were resuspended in formaldehyde lysis buffer (FALB; 50 mM HEPES pH 7.5, 140 mM NaCl, 1 mM EDTA, 1% Triton X-100) supplemented with 1% SDS and 1X cOmplete, EDTA-free Protease Inhibitor Cocktail (Roche cat. no. COEDTAF-RO), then incubated on ice for 15 minutes and sonicated for 21 minutes (10 seconds on/10 seconds off) at 25% power using an Active Motif Epishear Probe Sonicator. To remove debris, samples were centrifuged at 21000 x g for 10 minutes at 4 °C and the supernatant transferred to a new tube.

For immunoprecipitation, 30 μl Protein A Dynabeads (Invitrogen #10002D) per sample were washed with FALB (no SDS) thrice and blocked using 10 μg BSA for 20 minutes at room temperature. Beads were then incubated with anti-H3K27ac (Abcam ab4729) for 6-8 hours at 4°C on a rotator, before being washed with FALB (no SDS) thrice. Sonicated chromatin was sampled for input (2%) prior to a 1:10 dilution in FALB (no SDS) and overnight incubation at 4 °C on a rotator with the antibody-conjugated beads. The following day, beads were washed once for 5 minutes at 4 °C on a rotator in each of the following ice-cold buffers: high-salt wash buffer (20 mM Tris pH 8.0, 500 mM NaCl, 2 mM EDTA, 0.1% SDS, 1% Triton X-100), lithium chloride wash buffer (10 mM Tris pH 8.0, 250 mM LiCl, 1 mM EDTA, 1% IGEPAL), and TE wash buffer (10 mM Tris pH 8.0, 1 mM EDTA, 50 mM NaCl).

Immunoprecipitated chromatin was then eluted for 1 hour at 65 °C in elution buffer (50 mM Tris pH 8, 1 mM EDTA, 1% SDS), with eluate then transferred to a new tube. Input and ChIP samples were supplemented with 200mM NaCl and 40 μg proteinase K (Macherey-Nagel #740506), incubated for 30 minutes at 55 °C and then overnight at 65°C. DNA was then purified using the Zymo ChIP DNA Clean & Concentrator kit (#D5205).

Library preparation on input and ChIP samples was performed using Diagenode Microplex v3 (#C05010001) and UDIs (Set 2: #C05010009), according to manufacturer’s instructions and without size-selection. Libraries were sequenced paired-end 75 bp on Element AVITI by the IGC at Salk.

#### CUT&RUN and ChIP-seq analysis

##### Alignment and processing

Reads were aligned to both the hg38 and the E. coli K12, MG1655 reference genome assemblies (CUT&RUN) or to the hg38 assembly only (ChIP-seq) using the BWA-MEM (*97*). Optical and PCR duplicates were removed from both alignments using Picard MarkDuplicates (https://broadinstitute.github.io/picard/ (98)). Reads with a MAPQ<30, improper pairs, and supplementary aligned reads were removed using samtools (*102*). BEDTools intersect was then used to remove blacklisted regions (*103*) from the hg38 aligned reads.

##### Peak calling

Peaks were called from the hg38 aligned reads using SICER2 (*104*) and default settings, with the IgG as a control for CUT&RUN or input as a control for ChIP-seq.

##### Differential analysis

Differential enrichment of H3K27me3 (CUT&RUN), H3K4me3 (CUT&RUN) and H3K27ac (ChIP-seq) between IMR90 ^E6E7^ and PC7 was performed using DiffBind (*105*) based on the SICER2 peaks.

Normalization was performed using DBA_NORM_TMM, with the *E. coli* aligned reads used to normalize CUT&RUN data. Differential peaks were determined using the DBA_EDGER method. Peaks were classified as significantly changed based on an FDR<0.01.

*BigWig normalization:* BigWig files were normalized based on scale-factor (derived from DiffBind normFactor) using DeepTools bamCoverage (*106*).

##### Figures

Plotting of Hi-C heatmaps, BigWig and Virtual4C tracks was done using CoolBox (*99*).

**Fig. S1.**
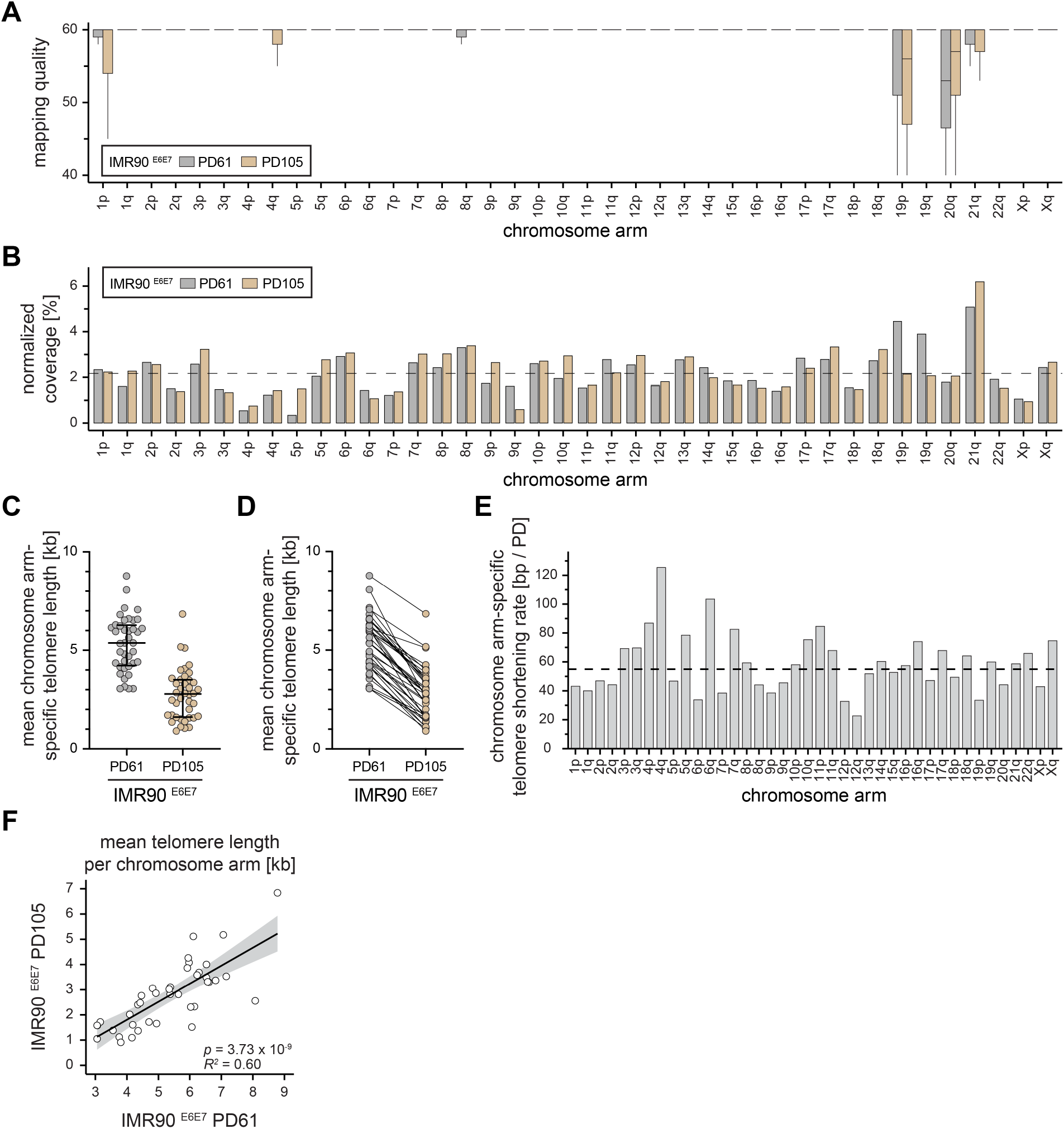
Telomere sequencing reveals short and heterogeneous telomere length in IMR90 E6E7 crisis cells. (**A**) Boxplot of telomere sequencing mapping quality per chromosome arm for IMR90 ^E6E7^ growing (PD61) and crisis (PD105) after filtering for reads with mapping quality of 40 and higher. The middle line represents median, the box the interquartile range (IQR) and whiskers the 1.5-fold IQR. The acrocentric p-arms of chromosomes 13, 14, 15, 21, and 22 and chromosome arm 20p could not be assembled in the custom reference genome and thus are missing. (**B**) Bar graph of the normalized coverage of mapped telomeric reads per chromosome arm for growing and crisis IMR90 ^E6E7^ cells in percent. Mapped 40+ telomeric reads per chromosome arm are normalized to total mapped 40+ telomeric reads. The expected average coverage for all chromosome arms in a diploid cell is shown as black dotted line. (C) Scatter plot of mean chromosome arm-specific telomere length in kilobases (kb). Median telomere length with IQR is shown. (**D**) Scatter of chromosome arm-specific mean telomere length in kb. Individual chromosome arms in growing and crisis IMR90 ^E6E7^ cells are linked by a line. (**E**) Bar graph of chromosome arm-specific telomere shortening rate in base pairs (bp) per population doubling (PD). The mean bulk telomere shortening rate is shown as black dotted line. (**F**) Linear regression analysis of chromosome arm-specific telomere length in kb for growing (PD61) and crisis (PD105) IMR90 ^E6E7^ cells. Black line represents best fit and the grey area 95% confidence interval.

**Fig. S2.**
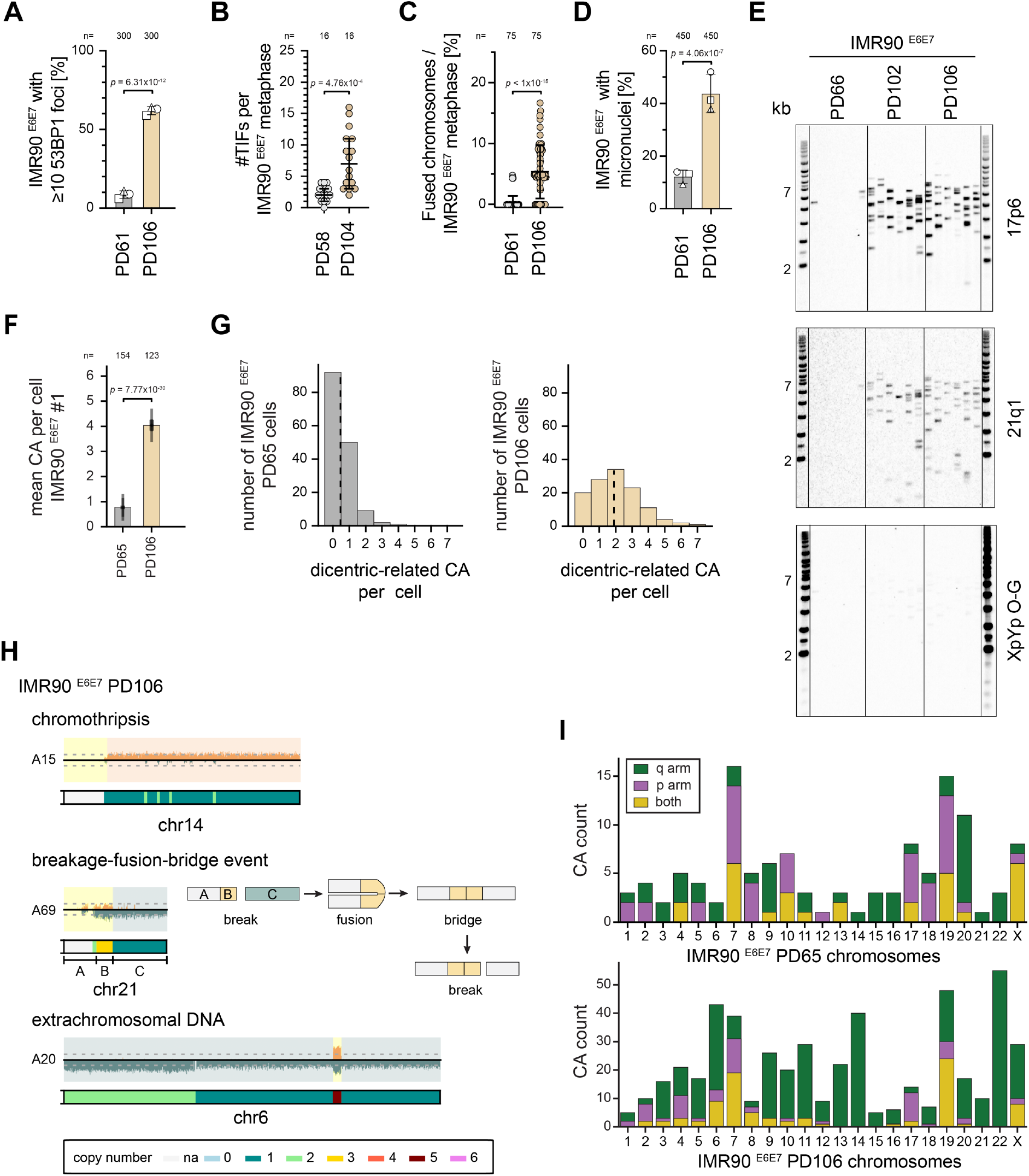
High and heterogeneous genomic instability in crisis. (**A**) Bar graph of percentage of growing (PD61) and crisis (PD106) IMR90 ^E6E7^ cells with at least 10 53BP1 foci. Mean and standard deviation of three independent experiments (100 cells per experiment) as well as the mean of the replicates are shown. Statistical test: Ordinary one-way ANOVA with Tukey’s multiple comparisons test. (**B**) Scatter plot of telomere dysfunction-induced foci (TIF) per metaphase for indicated cells. Individual TIFs per metaphase, the median with 95% confidence interval of two independent experiments with 8 metaphases each are shown. Statistical test: Kruskal-Wallis test with Dunn’s multiple comparison correction. (**C**) Scatter dot plot of the percentage of fused chromosomes per metaphase with mean and standard deviation. Each dot represents one metaphase. Cumulative data of three independent experiments with 25 metaphase each. Statistical test: Kruskal-Wallis test with Dunn’s multiple comparison correction compared to IMR90 ^E6E7^ PD61 growing control. (**D**) Bar graph of percentage of cells with micronuclei. Mean and standard deviation of three independent experiments (150 cells per experiment) as well as the mean of the replicates are shown. Statistical test: Ordinary one-way ANOVA with Tukey’s multiple comparisons test. (**E**) Telomeric fusion PCR assay for the indicative primer pairs and samples. Six independent reactions per sample. One experiment. (**F**) Bar graph of mean chromosomal abnormalities (CA) with standard deviation per cell in IMR90 ^E6E7^ growing and crisis cells. Population doubling (PD) is given. Statistical test: Mann-Whitney U test. (**G**) Histogram of dicentric-related CA per cell in IMR90 ^E6E7^ PD61 and PD106. Mean dicentric-related CA per cell is shown as vertical dashed line. (**H**) Representative examples of chromothripsis, breakage-fusion-bridge (BFB) events and extrachromosomal DNA detected in IMR90 ^E6E7^ PD106 Strand-seq. Upper panel: Strand-seq read distribution along chromosome coordinates, with Watson-oriented reads shown in orange and Crick-oriented reads in teal. Lower panel: Raw copy-number predictions generated by strandtools. Top: Chromothripsis is characterized by oscillations in copy-number states in the same and, less frequently, opposite strand orientations. Mid: BFB events display a stepwise increase in copy number extending from the centromere toward the chromosome end, often ending with an abrupt decrease. A schematic possible order of events is shown on the right. Bottom: A pattern consistent with extrachromosomal DNA (ecDNA) formation is characterized by copy-number loss across most of the chromosome, except for a relatively small segment that is highly amplified in both strand orientations. (**I**) Bar graph of CA per chromosome arm in growing (PD65) and crisis (PD106) IMR90 ^E6E7^ cells.

**Fig. S3.**
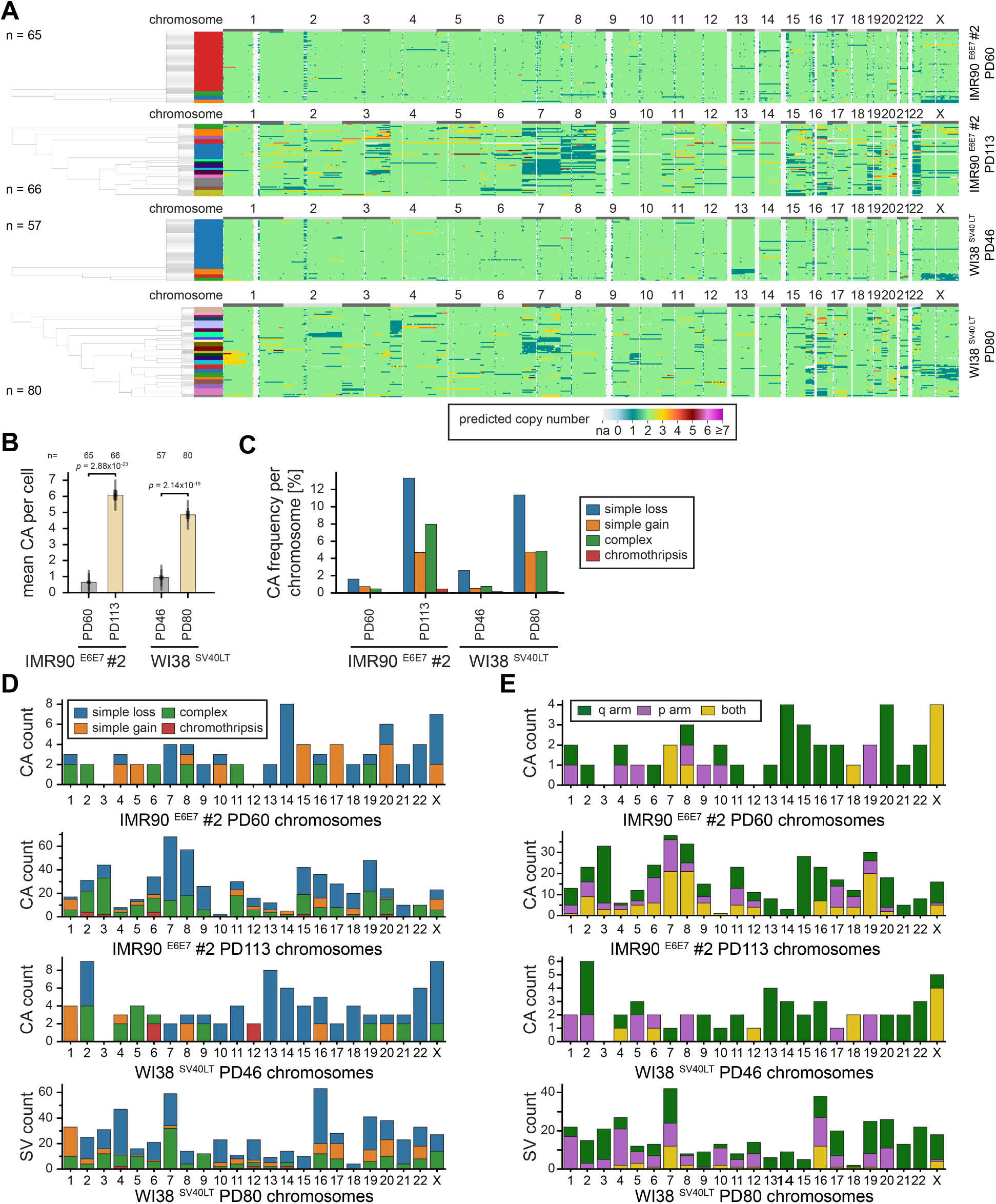
Single-cell analysis revealed high and heterogeneous genomic instability in crisis. (**A**) Copy number estimates of individual IMR90 ^E6E7^ #2 (second replicate independently transduced with HPV E6 and E7) growing (PD60) and crisis (PD113) cells and WI38 SV40LT growing (PD46) and crisis (PD80) cells based on single-cell Strand-seq. Phylogenetic trees were calculated with PICASSO and the most representative tree of eight PICASSO runs is shown. (**B**) Bar graph of mean chromosomal abnormalities (CA) with standard deviation per cell in growing and crisis cells. Population doubling (PD) is given. Statistical test: Mann-Whitney U test. (**C**) Bar graph of CA frequency per chromosome in percent of growing (PD60/46) and crisis (PD113/80) IMR90 ^E6E7^ #2 and WI38^SV40LT^ cells. (**D**) Bar graph of CA per chromosome in growing (PD60/46) and crisis (PD113/80) IMR90 ^E6E7^ #2 and WI38^SV40LT^ cells. (**E**) Bar graph of CA per chromosome arm in growing (PD60/46) and crisis (PD113/80) IMR90 ^E6E7^ #2 and WI38^SV40LT^ cells.

**Fig. S4.**
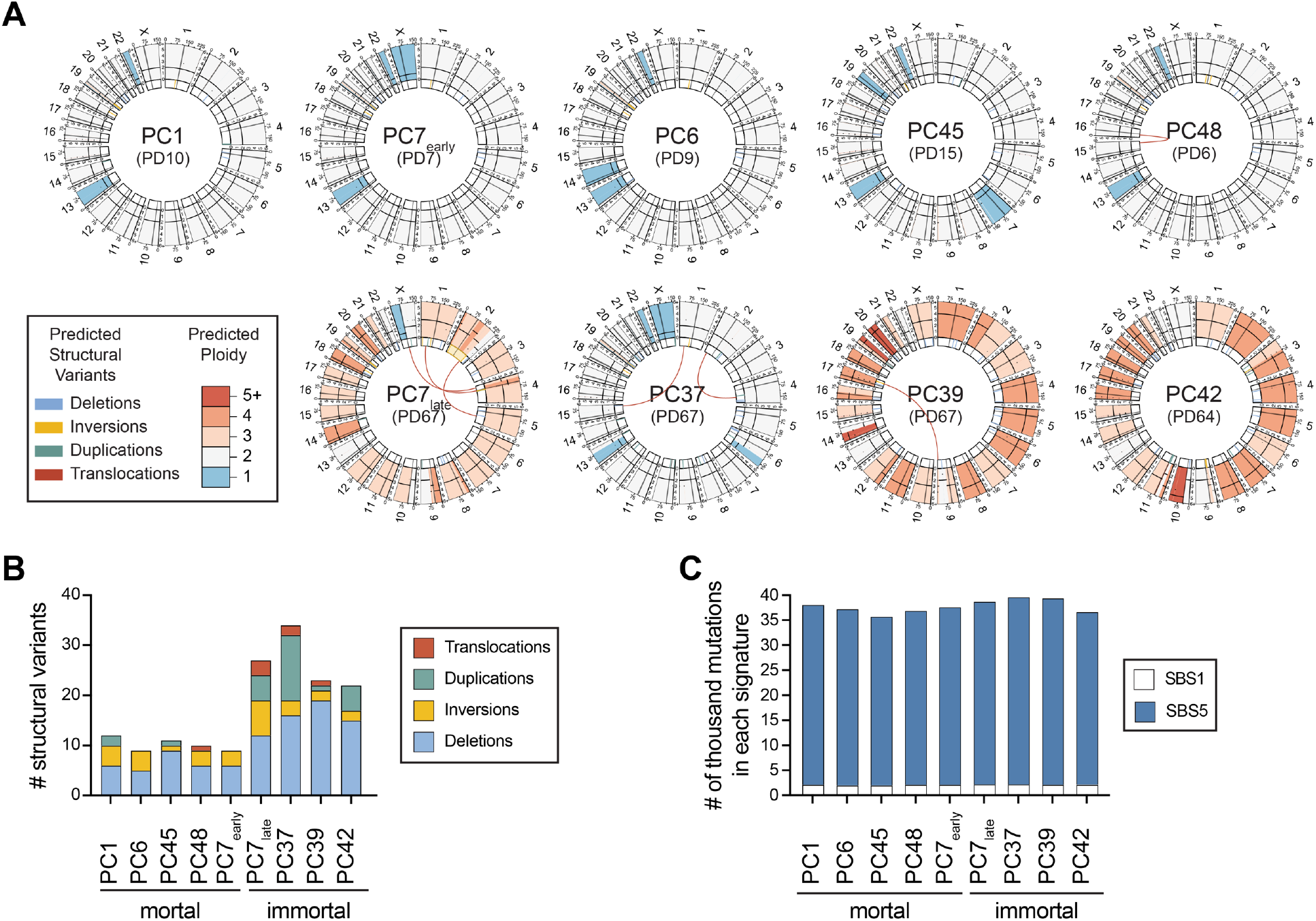
Postcrisis clones acquired relatively simple structural variants. (**A**) Circos plots of next-generation bulk whole genome sequencing of the indicated postcrisis clones at indicated population doubling (PD). (**B**) Bar graph of identified structural variants in bulk whole genome sequencing of postcrisis clones. (**C**) Stacked bar graph of SigProfilerAssignment single-base substitution mutational signature analysis of indicated postcrisis clones. SBS1 and SBS5 mutational signatures represent “spontaneous deamination of 5-hydroxycytosine (clock-like signature)” and “unknown (clock-like signature)”, respectively.

**Fig. S5.**
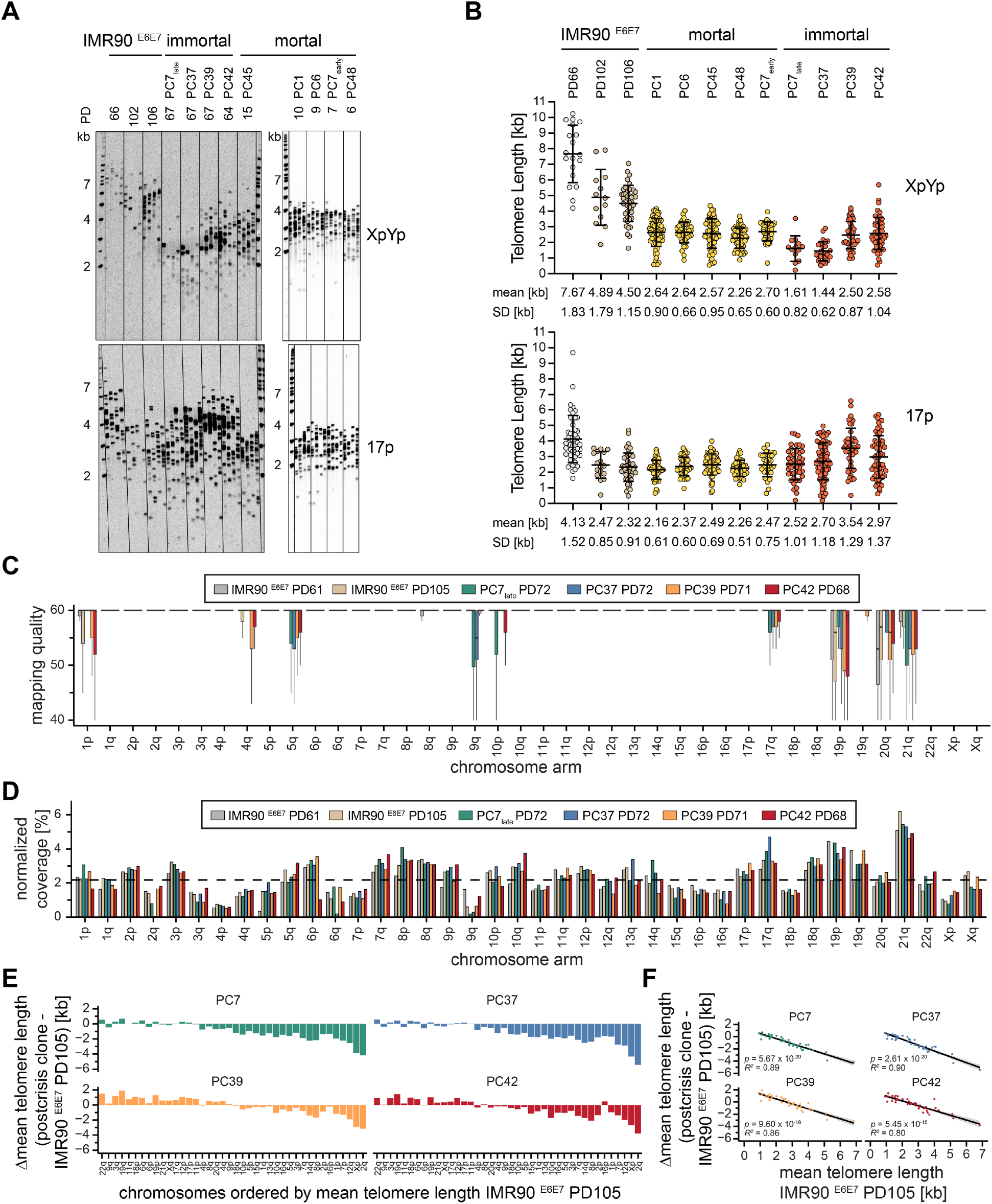
Postcrisis clones have short telomeres. (**A**) Single Telomere Length Analysis (STELA) analysis of IMR90 ^E6E7^ growing (population doubling (PD) 66), precrisis (PD102) and crisis (PD106) and postcrisis clones at indicated PDs. Four independent reactions were performed per sample. One experiment. (**B**) Quantification of STELA telomere length in kilobase (kb) shown in A. Mean and standard deviation (SD) are shown. (**C**) Boxplot of mapping quality per chromosome arm for indicated samples after filtering for reads with mapping quality of 40 and higher. The middle line represents median, the box the interquartile range (IQR) and whiskers the 1.5-fold IQR. The acrocentric p-arms of chromosomes 13, 14, 15, 21, and 22 and chromosome arm 20p could not be assembled in the used custom reference genome and thus are not shown. Growing (PD61) and crisis (PD105) IMR90 ^E6E7^ data is shown as reference and previously reported in fig. S1A. (**D**) Bar graph of the normalized coverage of mapped telomeric reads per chromosome arm for the indicated samples in percent. Mapped 40+ telomeric reads per chromosome arm are normalized to total mapped 40+ telomeric reads. The expected average coverage for all chromosome arms in a diploid cell is shown as black dotted line. Growing (PD61) and crisis (PD105) IMR90 ^E6E7^ data is shown as reference and previously reported in fig. S1B. (**E**) Bar graph of the postcrisis clone to IMR90 ^E6E7^ PD105 (crisis) mean chromosome arm-specific telomere length difference in kb. Chromosome arms are ordered according to mean telomere length in IMR90 ^E6E7^ PD105. (**F**) Linear regression analysis of the postcrisis clone to IMR90 ^E6E7^ PD105 (crisis) mean chromosome arm-specific telomere length difference in kb and mean chromosome arm-specific telomere length in kb for crisis IMR90 ^E6E7^ cells. The black line represents best fit and the grey area 95% confidence interval.

**Fig. S6.**
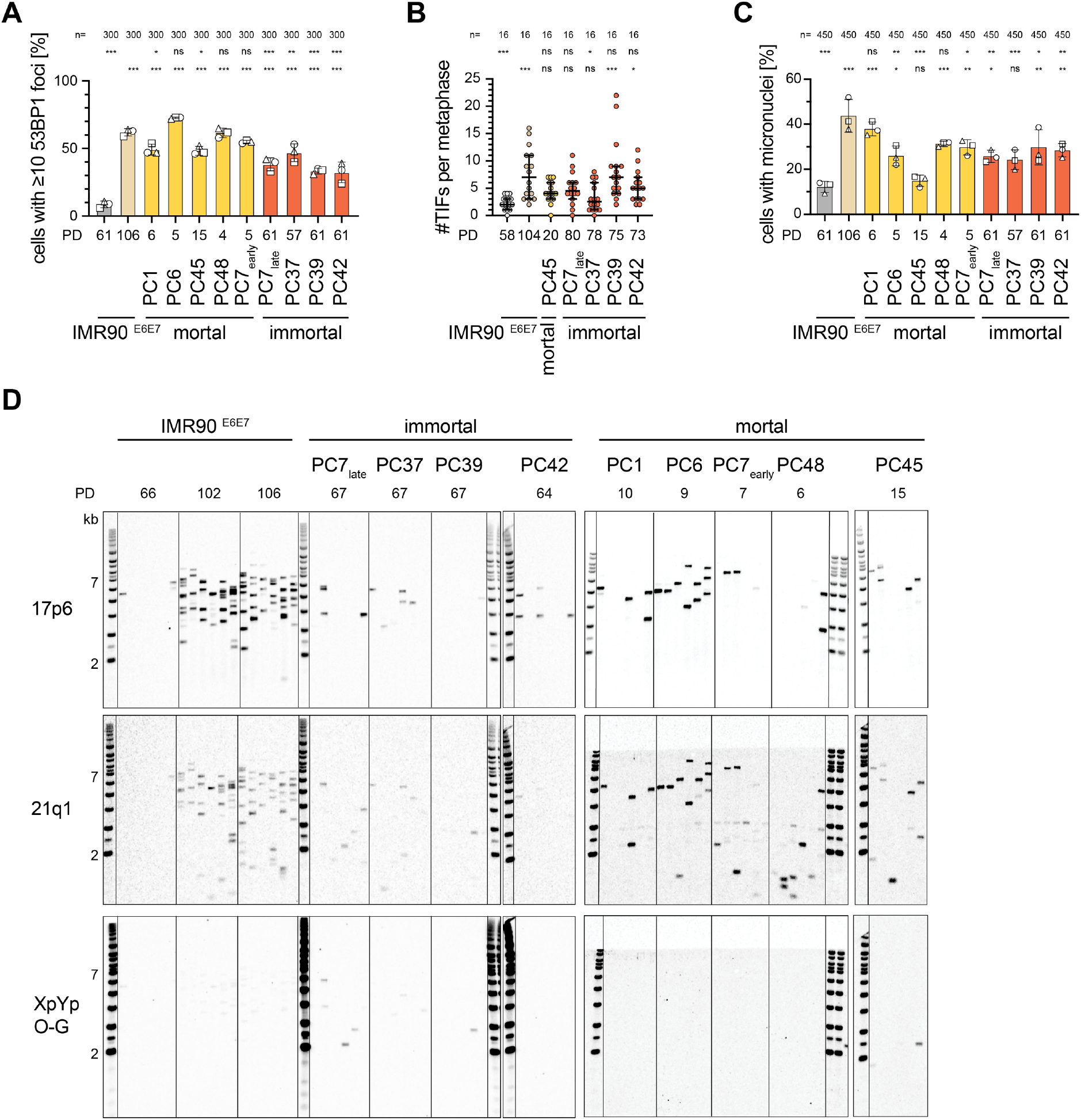
Postcrisis clones experience ongoing genome instability. (**A**) Bar graph of percentage of cells with at least 10 53BP1 foci. Mean and standard deviation of three independent experiments (100 cells per experiment) as well as the mean of the replicates are shown. Growing (PD61) and crisis (PD106) IMR90 ^E6E7^ data is shown as reference and previously reported in fig. S2A. Statistical test: Ordinary one-way ANOVA with Tukey’s multiple comparisons test. Adjusted p values correspond to * < 0.05, ** < 0.01, *** < 0.001 and ns for non-significant. Adjusted p values in comparison to IMR90 ^E6E7^ PD61 are: PD106 6.31×10^-12^, PC1 1.29×10^-9^, PC6 1.52×10^-13^, PC45 2.09×10^-9^, PC48 7.14×10^-12^, PC7early 1.00×10^-10^, PC7late 4.20×10^-7^, PC37 4.01×10^-9^, PC39 8.90×10^-6^, PC42 1.69×10^-5^. Adjusted p values in comparison to IMR90 ^E6E7^ PD106 are: PD61 6.31×10^-12^, PC1 0.0298, PC6 0.1036 PC45 0.0154, PC48 >0.9999, PC7early 0.5829, PC7late 1.69×10^-5^, PC37 6.19×10^-3^, PC39 7.56×10^-7^, PC42 4.20×10^-7^. (**B**) Scatter plot of telomere dysfunction-induced foci (TIF) per metaphase. Individual TIFs per metaphase, the median with 95% confidence interval of two independent experiments with 8 metaphases each are shown. Growing (PD58) and crisis (PD104) IMR90 ^E6E7^ data is shown as reference and previously reported in fig. S2B. Statistical test Kruskal-Wallis test with Dunn’s multiple comparison correction. * p < 0.05, ** p < 0.01, *** p < 0.001, ns for non-significant. Adjusted p values in comparison to IMR90 ^E6E7^ PD58 are: PD104 4.76×10^-4^, PC45 0.2923, PC7late 0.1050, PC37 >0.9999, PC39 9.81×10^-5^, PC42 0.0425. Adjusted p values in comparison to IMR90 ^E6E7^ PD106 are: PD61 4.76×10^-4^, PC45 >0.9999, PC7late >0.9999, PC37 0.0258, PC39 >0.9999, PC42 >0.9999. (**C**) Bar graph of percentage of cells with micronuclei. Mean and standard deviation of three independent experiments (150 cells per experiment) as well as the mean of the replicates are shown. Growing (PD61) and crisis (PD106) IMR90 ^E6E7^ data is shown as reference and previously reported in fig. S2D. Statistical test: Ordinary one-way ANOVA with Tukey’s multiple comparisons test. Adjusted p values correspond to * < 0.05, ** < 0.01, *** < 0.001 and ns for non-significant. Adjusted p values in comparison to IMR90 ^E6E7^ PD61 are: PD106 4.06×10^-7^, PC1 1.15×10^-5^, PC6 0.0241, PC45 0.9992, PC48 7.66×10^-4^, PC7early 2.09×10^-3^, PC7late 0.0278, PC37 0.0696, PC39 2.09×10^-3^, PC42 5.09×10^-3^. Adjusted p values in comparison to IMR90 ^E6E7^ PD106 are: PD61 4.06×10^-7^, PC1 0.8461, PC6 1.87×10^-3^, PC45 1.85×10^-6^, PC48 0.0551, PC7early 0.0217, PC7late 1.61×10^-3^, PC37 5.90×10^-4^, PC39 0.0217, PC42 9.11×10^-3^. (**D**) Telomeric fusion PCR assay for the indicative primer pairs and samples. Growing (PD66) and crisis (PD102 and PD106) IMR90 ^E6E7^ data is shown as reference and previously reported in fig. S2E. Six independent reactions per sample. One experiment.

**Fig. S7.**
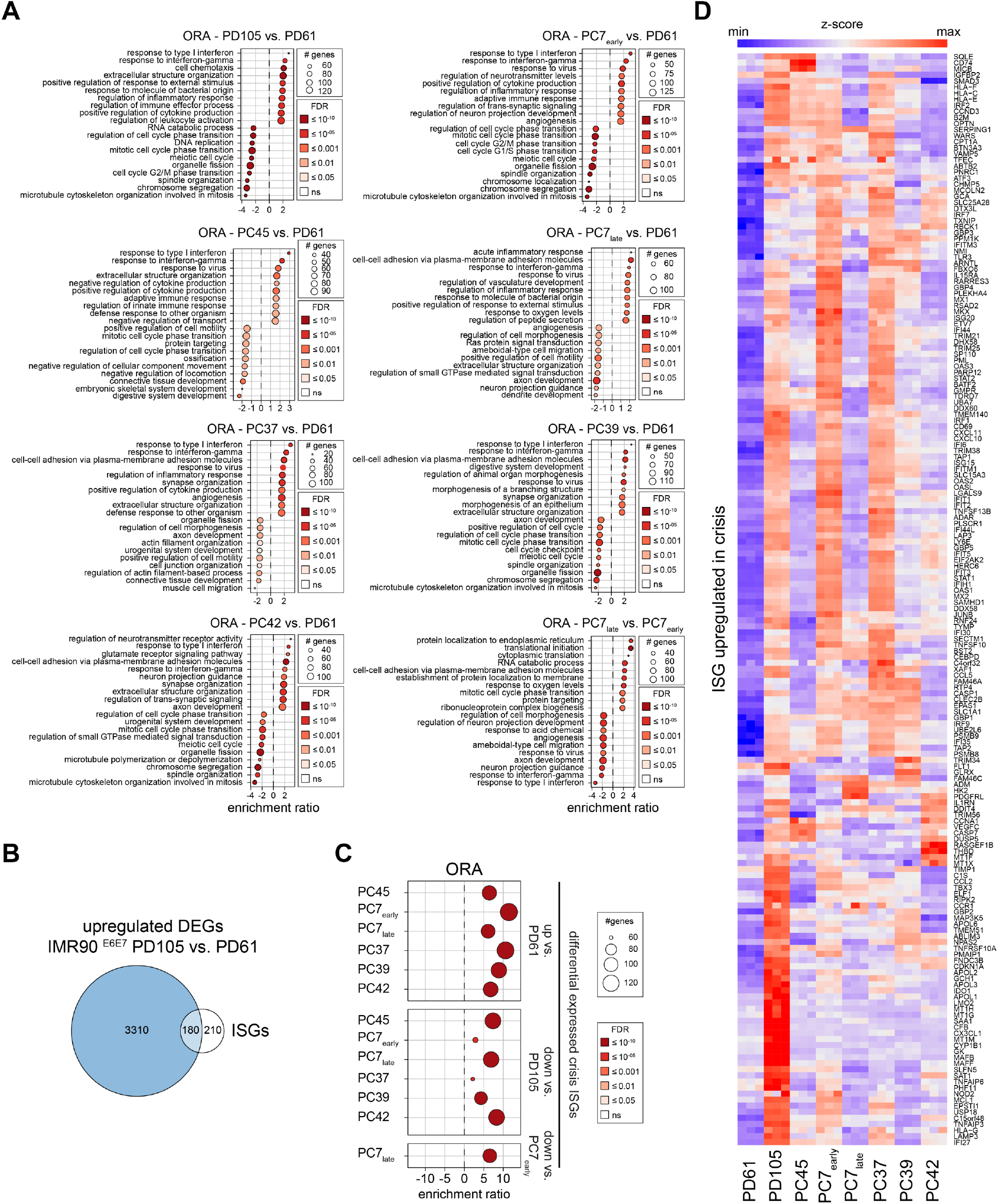
Immortal postcrisis clones express intermediate levels of interferon-stimulated genes. (**A**) Overrepresentation analysis (ORA) bulk RNA-sequencing expression data of indicated samples relative to growing IMR90 ^E6E7^ (population doubling (PD) 61) controls. (**B**) Venn diagram of upregulated, differential expressed genes (DEG) in IMR90 ^E6E7^ crisis (PD105) to growing (PD61) cells to list of Interferon-stimulated genes (ISGs) (*107*). (**C**) ORA of ISGs that were differentially upregulated in crisis (PD105) relative to growing (PD61) IMR90 ^E6E7^ cells for the indicated comparisons. (**D**) Heatmap of ISGs significantly upregulated in crisis. A-D are based on bulk RNA-sequencing with three replicates per sample. Bulk RNA-sequencing was performed at the following PDs: IMR90 ^E6E7^ PD61 and PD105, PC45 PD19, PC7early PD5, PC7late PD59, PC37 PD58, PC39 PD54, and PC42 PD68.

**Fig. S8.**
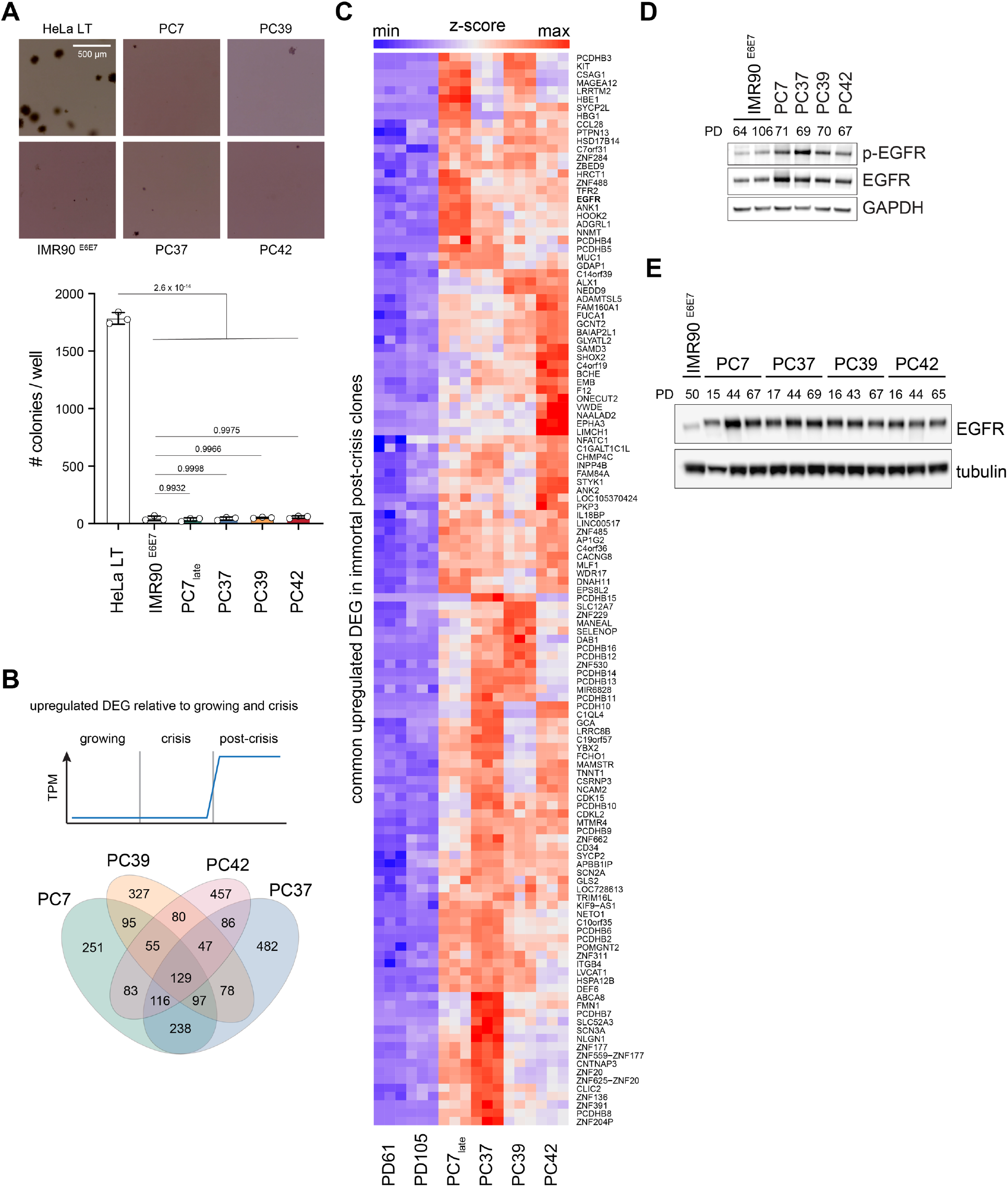
EGFR is overexpressed in immortal postcrisis cells. (**A**) Soft agar assay of the indicated samples. HeLa LT cancer cell lines are used as positive controls. Representative of three independent experiments with three technical replicates is shown. Soft agar assay was performed at the following PDs: IMR90 ^E6E7^ PD70, PC7 PD73, PC37 PD72, PC39 PD73 and PC42 PD69. Statistical test: Ordinary one-way ANOVA with Tukey’s multiple comparisons test. Adjusted p values are shown. (**B**) Venn diagram of differentially upregulated genes in RNA-sequencing data of immortal postcrisis clones relative to growing and crisis IMR90 ^E6E7^ cells. In the bulk RNA-sequencing three replicates per sample were analyzed. Bulk RNA-sequencing was performed at the following population doublings (PD): IMR90 ^E6E7^ PD61 and PD105, PC7late PD59, PC37 PD58, PC39 PD54, and PC42 PD68. (**C**) Heat map of the 129 differentially upregulated genes shared between all immortal postcrisis clones in bulk RNA-sequencing. (**D**) Western blot analysis of total and Y1068 phosphorylated EGFR protein levels in the indicated cells. Three independent experiments. (**E**) Western blot analysis of EGFR protein levels in immortal postcrisis clones at different PDs. Three independent experiments.

**Fig. S9.**
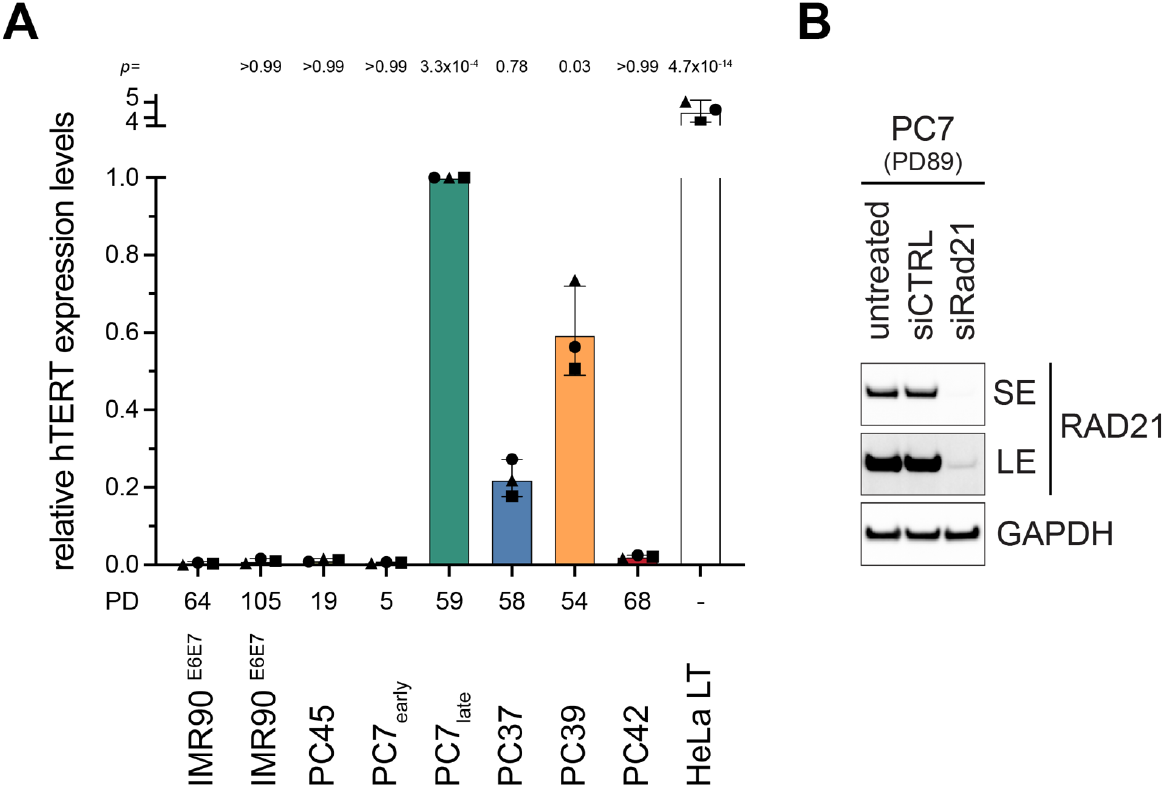
Immortal postcrisis clones express TERT. (**A**) Bar graph of relative hTERT expression levels measured by RT-qPCR and relative to PC7late, the postcrisis clone with highest TERT expression. Mean of three independent experiments with standard deviation and the mean for each individual experiment are shown. Per individual experiment three technical replicates were measured. (**B**) Western blot analysis of Rad21 protein levels in untransfected, control siRNA (siCTRL) or siRNA targeting Rad21 (siRad21) transfected PC7 cells. For Rad21 a short (SE) and long (LE) exposure is shown. Three independent experiments.

**Fig. S10.**
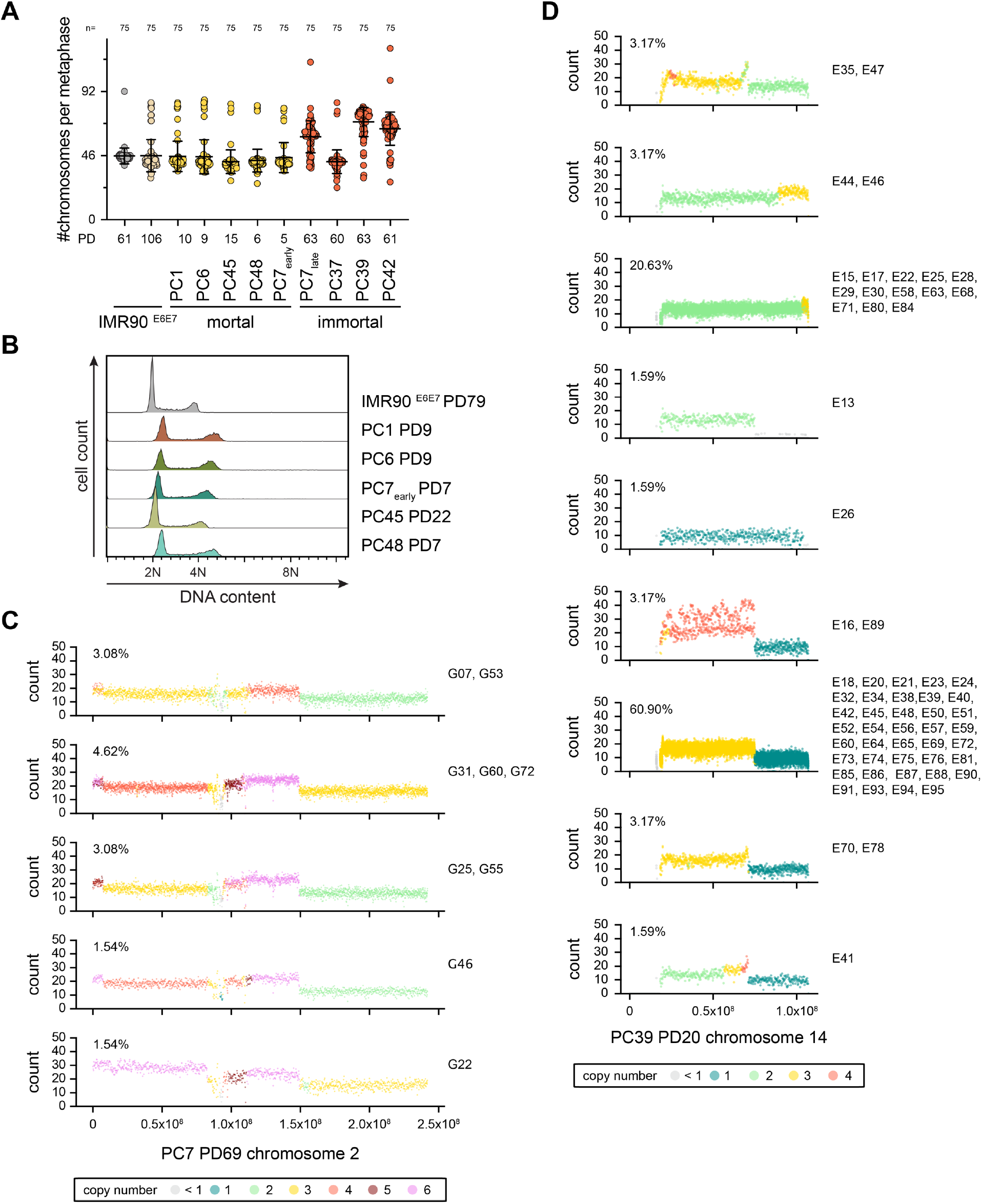
Postcrisis clones escape crisis with near diploid ploidy. (**A**) Scatter dot plot of chromosomes per metaphase quantified by centromeric FISH staining with mean and standard deviation. Each dot represents one metaphase. Cumulative data of three independent experiments with 25 metaphase each. (**B**) DNA content analysis of indicated samples. IMR90 ^E6E7^ (population doubling (PD) 79) represent parental diploid control cells. Three independent experiments. (**C-D**) Subclonal read-depth plots from Strand-seq data for the events indicative of breakage-fusion-bridge (BFB) events in the immortal postcrisis clones. Cells were hierarchically clustered on the basis of the per-bin predicted copy number states across the chromosome of interest, with Ward linkage and Euclidean distance. The dendrogram was cut at a series of distance thresholds to generate increasing numbers of clusters, and the optimal number of clusters was selected by visual inspection of the elbow in this relationship. For each cluster, total read counts at each genomic bin were then plotted across cells to generate consensus subclone-level copy number profiles. Dots are colored by predicted copy number state. Cell IDs and frequency are shown. BFB events display a stepwise increase in copy number extending from the centromere toward the chromosome end, often ending with an abrupt decrease.

**Fig. S11.**
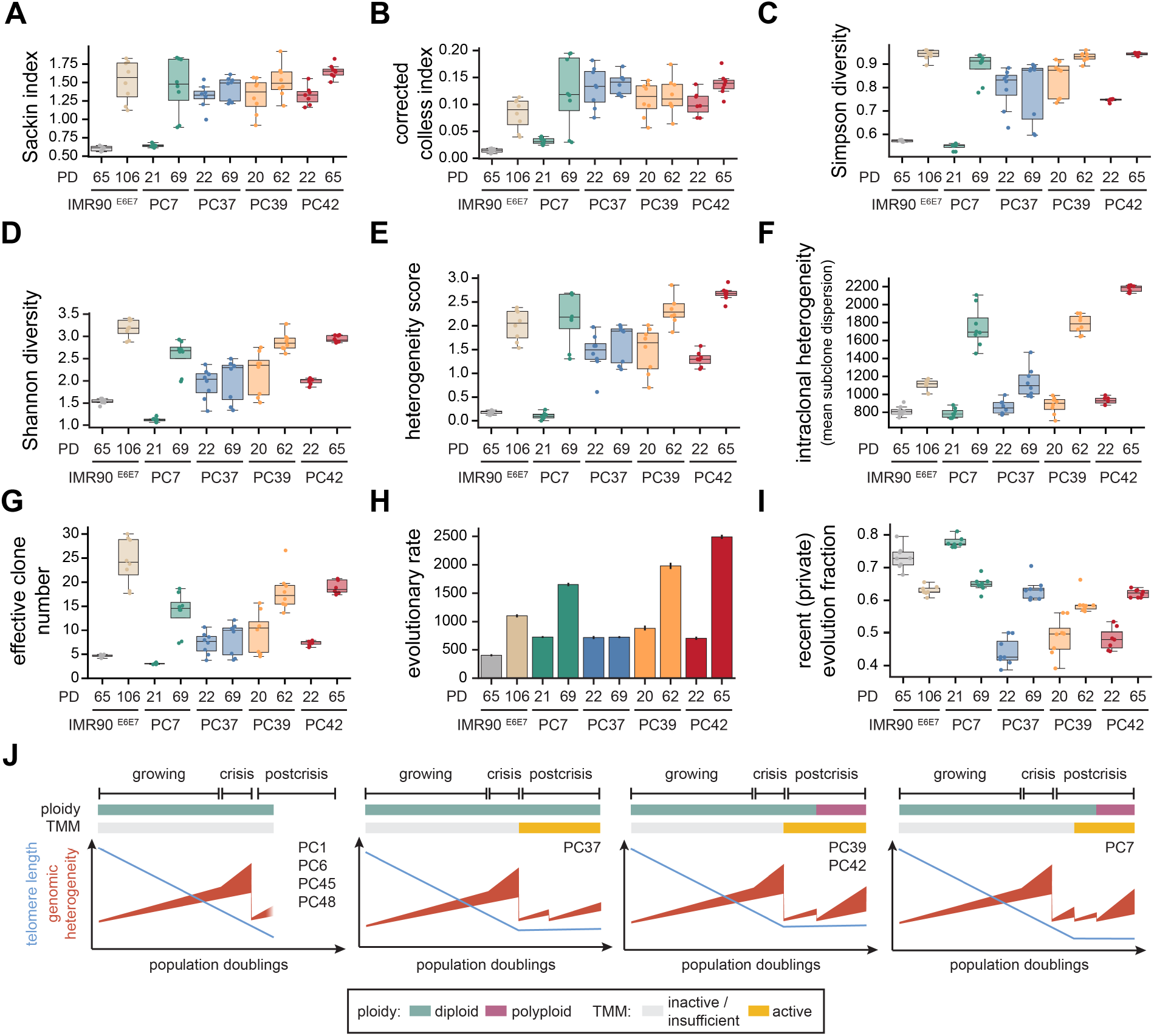
Immortal postcrisis clones are evolving. (**A-I**) Indicated phylogenetic tree statistic parameter based on PICASSO analysis of copy number estimates of single-cell Strand-seq of IMR90 ^E6E7^ growing (PD65), crisis (PD106) and immortal postcrisis clones of the indicated PD. PICASSO was run eight times. For Boxplots the middle line represents median, the box the interquartile range (IQR) and whiskers the 1.5-fold IQR. Bar graph shown in (H) is the mean evolutionary rate with 95% confidence intervals. For more information, see Materials and Methods. (**J**) Model for evolution of cells through crisis to postcrisis. Different trajectories exemplified by different postcrisis clones are shown.

**Table S1.** Bulk telomere length of IMR90 ^E6E7^ growing, crisis and immortal postcrisis clones.

|  | IMR90<sup>E6E7</sup> |  | immortal postcrisis clone |  |  |  |
| --- | --- | --- | --- | --- | --- | --- |
| Sample | growing | crisis | PC7 | PC37 | PC39 | PC42 |
| Population doublings | 61 | 105 | 72 | 72 | 71 | 68 |
| **Flow cell** |  |  |  |  |  |  |
| Flow cell ID | PAW27871 |  |  |  |  |  |
| No. total reads | 2220864 |  |  |  |  |  |
| No. all non-telomeric reads | 2020197 |  |  |  |  |  |
| No. all telomeric reads | 200667 |  |  |  |  |  |
| % telomeric / total reads | (9.03%) |  |  |  |  |  |
| Telomeric adapter | TA1 | TA2 | TA3 | TA4 | TA5 | TA6 |
| No. of telomeric reads / sample | 24453 | 35185 | 30810 | 24435 | 36305 | 49479 |
| No. of mapped telomeric reads Q40+ | 18507 | 28130 | 24684 | 19625 | 28510 | 39630 |
| % mapped reads Q40+ | 75.68 | 79.95 | 80.12 | 80.32 | 78.53 | 80.09 |
| **Telomere length** |  |  |  |  |  |  |
| Mean [bp] | 5139 | 2721 | 1797 | 1588 | 2631 | 2297 |
| Standard deviation [bp] | 2741 | 2002 | 805 | 746 | 1166 | 1143 |
| Min [bp] | 67 | 73 | 90 | 105 | 74 | 97 |
| 10<sup>th</sup> percentile [bp] | 2049 | 803 | 855 | 709 | 1235 | 981 |
| 25<sup>th</sup> percentile [bp] | 3155 | 1326 | 1218 | 1041 | 1782 | 1466 |
| Median [bp] | 4740 | 2225 | 1696 | 1489 | 2500 | 2142 |
| 75<sup>th</sup> percentile [bp] | 6660 | 3514 | 2268 | 2025 | 3334 | 2951 |
| 90<sup>th</sup> percentile [bp] | 8640 | 5344 | 2868 | 2607 | 4184 | 3799 |
| Max [bp] | 39517 | 39506 | 10356 | 6333 | 8998 | 15386 |
| IQR [bp] | 3505 | 2188 | 1050 | 984 | 1552 | 1485 |
| CV | 0.53 | 0.74 | 0.45 | 0.47 | 0.44 | 0.50 |
| % telomeres <1 kb | 1.71 | 15.45 | 15.10 | 22.86 | 5.73 | 10.40 |
| % telomeres >10 kb | 4.96 | 0.84 | 0.003 | 0 | 0 | 0.03 |

**Table S2.** Primary antibody list.

| target | species | clone | supplier | catalog number | RRID | assay | dilution |
| --- | --- | --- | --- | --- | --- | --- | --- |
| 53BP1 | rabbit | polyclonal | Santa Cruz Biotechnology | sc-22760 | AB_2256326 | IF | 1:2000 |
| EGFR | rabbit | polyclonal | CST | 4267 | AB_2246311 | WB | 1:1000 |
| phospho-EGFR (TYR1068) | rabbit | D7A5 | CST | 3777 | AB_2096270 | WB | 1:1000 |
| GAPDH | mouse | polyclonal | Abnova | PAB17013 | AB_10629928 | WB | 1:5000 |
| GAPDH | rabbit | 14C10 | CST | 2118 | AB_561053 | WB | 1:1000 |
| phospho-H2A.X (Ser139) | mouse | JBW301 | Millipore | 05-636 | AB_309864 | IF | 1:1000 |
| IFIT1 | rabbit | D2X9Z | CST | 14769 | AB_2783869 | WB | 1:1000 |
| ISG15 | rabbit | 22D2 | CST | 2758 | AB_2126200 | WB | 1:1000 |
| LC3B | rabbit | polyclonal | CST | 2775 | AB_915950 | WB | 1:1000 |
| RAD21 | rabbit | polyclonal | Bethyl | A300-080A | AB_2176615 | WB | 1:25000 |
| RIG1 | rabbit | D14G6 | CST | 3743 | AB_2269233 | WB | 1:1000 |
| phospho-STAT1 (Tyr701) | rabbit | 58D6 | CST | 9167 | AB_561284 | WB | 1:1000 |
| TBK1/NAK | rabbit | D1B4 | CST | 3504 | AB_2255663 | WB | 1:1000 |
| phospho-TBK1 (Ser172) | rabbit | D52C2 | CST | 5483 | AB_10693472 | WB | 1:1000 |
| γ-tubulin | mouse | GTU-88 | Sigma | T6557 | AB_477584 | WB | 1:5000 |

**Table S3.**
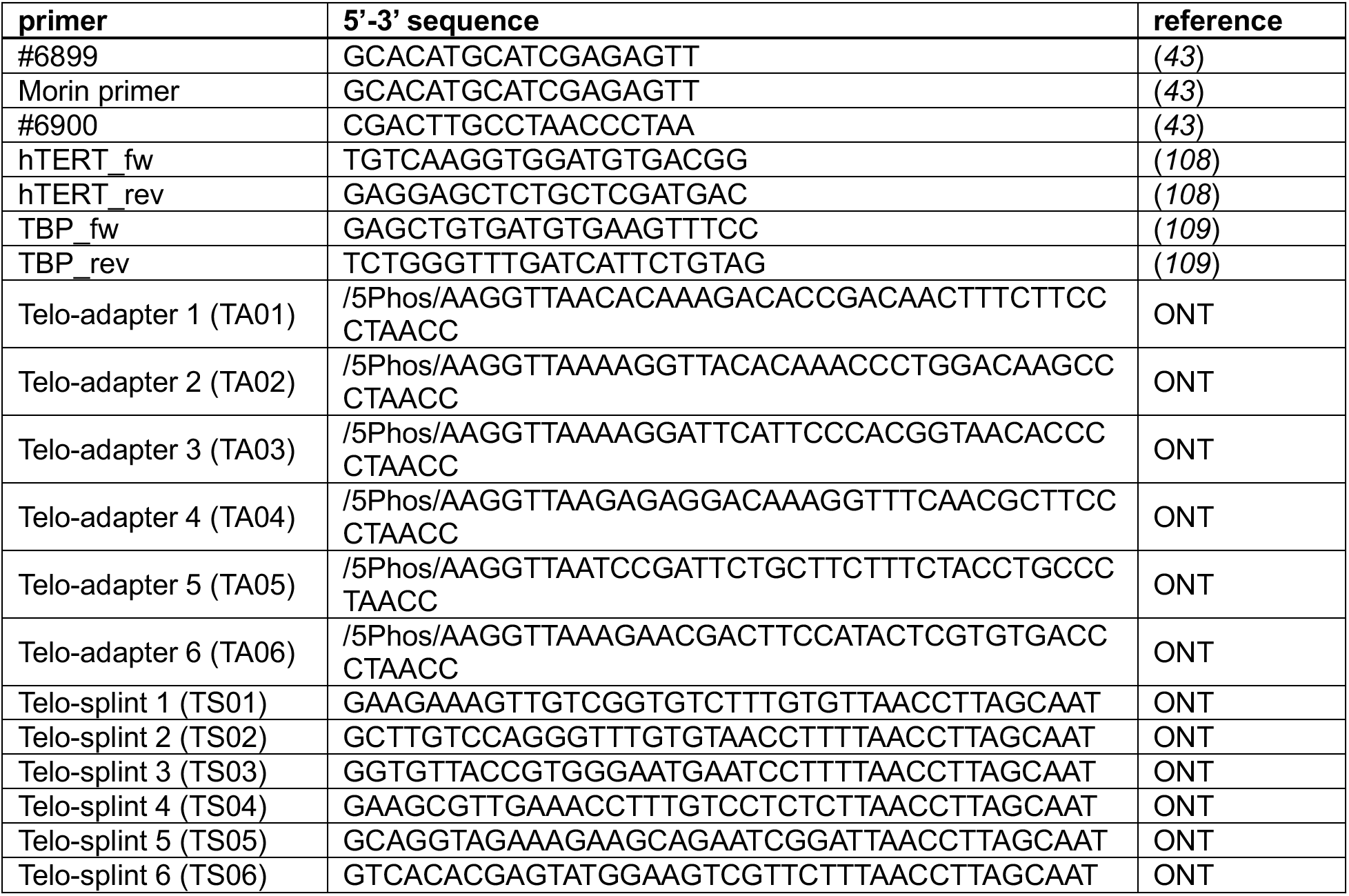
Oligos used in this study.

**Data S1. Chromosome arm-specific telomere length of IMR90 ^E6E7^ growing, crisis and immortal postcrisis clones.**

## References and Notes

1. D. Hanahan, Hallmarks of cancer-Then and now, and beyond. Cell 189, 2254–2277 (2026).

2. T. Baslan et al., Ordered and deterministic cancer genome evolution after p53 loss. Nature 608, 795–802 (2022).

3. S. F. Bakhoum et al., Chromosomal instability drives metastasis through a cytosolic DNA response. Nature 553, 467–472 (2018).

4. J. Li et al., Non-cell-autonomous cancer progression from chromosomal instability. Nature 620, 1080–1088 (2023).

5. T. B. K. Watkins et al., Pervasive chromosomal instability and karyotype order in tumour evolution. Nature 587, 126–132 (2020).

6. L. Sansregret, B. Vanhaesebroeck, C. Swanton, Determinants and clinical implications of chromosomal instability in cancer. Nat Rev Clin Oncol 15, 139–150 (2018).

7. M. R. Cosenza, B. Rodriguez-Martin, J. O. Korbel, Structural Variation in Cancer: Role, Prevalence, and Mechanisms. Annu Rev Genomics Hum Genet 23, 123–152 (2022).

8. M. Meyerson et al., hEST2, the putative human telomerase catalytic subunit gene, is up-regulated in tumor cells and during immortalization. Cell 90, 785–795 (1997).

9. T. M. Bryan, A. Englezou, L. Dalla-Pozza, M. A. Dunham, R. R. Reddel, Evidence for an alternative mechanism for maintaining telomere length in human tumors and tumor-derived cell lines. Nat Med 3, 1271–1274 (1997).

10. J. D. Watson, Origin of concatemeric T7 DNA. Nat New Biol 239, 197–201 (1972).

11. A. M. Olovnikov, A theory of marginotomy. The incomplete copying of template margin in enzymic synthesis of polynucleotides and biological significance of the phenomenon. J Theor Biol 41, 181–190 (1973).

12. J. Lingner, J. P. Cooper, T. R. Cech, Telomerase and DNA end replication: no longer a lagging strand problem? Science 269, 1533–1534 (1995).

13. L. Hayflick, The Limited in Vitro Lifetime of Human Diploid Cell Strains. Exp Cell Res 37, 614–636 (1965).

14. J. W. Shay, O. M. Pereira-Smith, W. E. Wright, A role for both RB and p53 in the regulation of human cellular senescence. Exp Cell Res 196, 33–39 (1991).

15. W. E. Wright, O. M. Pereira-Smith, J. W. Shay, Reversible cellular senescence: implications for immortalization of normal human diploid fibroblasts. Mol Cell Biol 9, 3088–3092 (1989).

16. G. H. Stein, SV40-transformed human fibroblasts: evidence for cellular aging in pre-crisis cells. J Cell Physiol 125, 36–44 (1985).

17. J. Nassour et al., Telomere-to-mitochondria signalling by ZBP1 mediates replicative crisis. Nature 614, 767–773 (2023).

18. J. Nassour et al., Autophagic cell death restricts chromosomal instability during replicative crisis. Nature 565, 659–663 (2019).

19. R. Capper et al., The nature of telomere fusion and a definition of the critical telomere length in human cells. Genes Dev 21, 2495–2508 (2007).

20. K. Liddiard, J. W. Grimstead, K. Cleal, A. Evans, D. M. Baird, Tracking telomere fusions through crisis reveals conflict between DNA transcription and the DNA damage response. NAR Cancer 3, zcaa044 (2021).

21. K. Liddiard et al., Sister chromatid telomere fusions, but not NHEJ-mediated inter-chromosomal telomere fusions, occur independently of DNA ligases 3 and 4. Genome Res 26, 588–600 (2016).

22. J. Nassour, T. T. Schmidt, J. Karlseder, Telomeres and Cancer: Resolving the Paradox. Annu Rev Cancer Biol 5, 59–77 (2021).

23. J. W. Shay, W. E. Wright, Quantitation of the frequency of immortalization of normal human diploid fibroblasts by SV40 large T-antigen. Exp Cell Res 184, 109–118 (1989).

24. F. A. Ray, P. M. Kraemer, Iterative chromosome mutation and selection as a mechanism of complete transformation of human diploid fibroblasts by SV40 T antigen. Carcinogenesis 14, 1511–1516 (1993).

25. C. Jones-Weinert, L. Mainz, J. Karlseder, Telomere function and regulation from mouse models to human ageing and disease. Nat Rev Mol Cell Biol 26, 297–313 (2025).

26. K. R. Prowse, C. W. Greider, Developmental and tissue-specific regulation of mouse telomerase and telomere length. Proc Natl Acad Sci U S A 92, 4818–4822 (1995).

27. R. Smoom, D. Lichtental, K. H. Kaestner, Y. Tzfati, The house mouse maintains constant telomere length throughout life. Nucleic Acids Res 53, (2025).

28. D. Kipling, H. J. Cooke, Hypervariable ultra-long telomeres in mice. Nature 347, 400–402 (1990).

29. M. A. Blasco et al., Telomere shortening and tumor formation by mouse cells lacking telomerase RNA. Cell 91, 25–34 (1997).

30. C. B. Harley, A. B. Futcher, C. W. Greider, Telomeres shorten during ageing of human fibroblasts. Nature 345, 458–460 (1990).

31. I. Cortes-Ciriano et al., Comprehensive analysis of chromothripsis in 2,658 human cancers using whole-genome sequencing. Nat Genet 52, 331–341 (2020).

32. T. T. Schmidt et al., High resolution long-read telomere sequencing reveals dynamic mechanisms in aging and cancer. Nat Commun 15, 5149 (2024).

33. A. D. Sanders et al., Single-cell analysis of structural variations and complex rearrangements with tri-channel processing. Nat Biotechnol 38, 343–354 (2020).

34. V. C. T. Hanlon et al., Construction of Strand-seq libraries in open nanoliter arrays. Cell Rep Methods 2, 100150 (2022).

35. M. R. Cosenza et al., Origins of chromosome instability unveiled by coupled imaging and genomics. Nature, (2025).

36. B. McClintock, The Stability of Broken Ends of Chromosomes in Zea Mays. Genetics 26, 234–282 (1941).

37. P. J. Stephens et al., Massive genomic rearrangement acquired in a single catastrophic event during cancer development. Cell 144, 27–40 (2011).

38. J. O. Korbel, P. J. Campbell, Criteria for inference of chromothripsis in cancer genomes. Cell 152, 1226–1236 (2013).

39. L. B. Alexandrov et al., The repertoire of mutational signatures in human cancer. Nature 578, 94–101 (2020).

40. D. M. Baird, J. Rowson, D. Wynford-Thomas, D. Kipling, Extensive allelic variation and ultrashort telomeres in senescent human cells. Nat Genet 33, 203–207 (2003).

41. V. H. Freedman, S. I. Shin, Cellular tumorigenicity in nude mice: correlation with cell growth in semi-solid medium. Cell 3, 355–359 (1974).

42. Y. Yarden, M. X. Sliwkowski, Untangling the ErbB signalling network. Nat Rev Mol Cell Biol 2, 127–137 (2001).

43. T. Rivera, C. Haggblom, S. Cosconati, J. Karlseder, A balance between elongation and trimming regulates telomere stability in stem cells. Nat Struct Mol Biol 24, 30–39 (2017).

44. S. Horn et al., TERT promoter mutations in familial and sporadic melanoma. Science 339, 959–961 (2013).

45. F. W. Huang et al., Highly recurrent TERT promoter mutations in human melanoma. Science 339, 957–959 (2013).

46. Y. Zhao, S. Wang, E. Y. Popova, S. A. Grigoryev, J. Zhu, Rearrangement of upstream sequences of the hTERT gene during cellular immortalization. Genes Chromosomes Cancer 48, 963–974 (2009).

47. M. Peifer et al., Telomerase activation by genomic rearrangements in high-risk neuroblastoma. Nature 526, 700–704 (2015).

48. L. J. Valentijn et al., TERT rearrangements are frequent in neuroblastoma and identify aggressive tumors. Nat Genet 47, 1411–1414 (2015).

49. Z. Xu et al., Structural variants drive context-dependent oncogene activation in cancer. Nature 612, 564–572 (2022).

50. A. Zhang et al., Frequent amplification of the telomerase reverse transcriptase gene in human tumors. Cancer Res 60, 6230–6235 (2000).

51. F. P. Barthel et al., Systematic analysis of telomere length and somatic alterations in 31 cancer types. Nat Genet 49, 349–357 (2017).

52. S. C. Akincilar et al., Long-Range Chromatin Interactions Drive Mutant TERT Promoter Activation. Cancer Discov 6, 1276–1291 (2016).

53. A. Sen et al., Heterogeneity in chromatin structure drives core regulatory pathways in B-cell Acute Lymphoblastic Leukemia. bioRxiv, 2024.2010.2004.616668 (2024).

54. D. D. Lee et al., DNA hypermethylation within TERT promoter upregulates TERT expression in cancer. J Clin Invest 129, 223–229 (2019).

55. I. Guilleret et al., Hypermethylation of the human telomerase catalytic subunit (hTERT) gene correlates with telomerase activity. Int J Cancer 101, 335–341 (2002).

56. R. L. Zinn, K. Pruitt, S. Eguchi, S. B. Baylin, J. G. Herman, hTERT is expressed in cancer cell lines despite promoter DNA methylation by preservation of unmethylated DNA and active chromatin around the transcription start site. Cancer Res 67, 194–201 (2007).

57. A. Jimenez-Sanchez et al., Transcriptomic Plasticity Is a Hallmark of Metastatic Pancreatic Cancer. Cancer Res 86, 1769–1796 (2026).

58. S. Gemble et al., Genetic instability from a single S phase after whole-genome duplication. Nature 604, 146–151 (2022).

59. R. A. Lambuta et al., Whole-genome doubling drives oncogenic loss of chromatin segregation. Nature 615, 925–933 (2023).

60. M. T. Hemann, M. A. Strong, L. Y. Hao, C. W. Greider, The shortest telomere, not average telomere length, is critical for cell viability and chromosome stability. Cell 107, 67–77 (2001).

61. Y. Zou, A. Sfeir, S. M. Gryaznov, J. W. Shay, W. E. Wright, Does a sentinel or a subset of short telomeres determine replicative senescence? Mol Biol Cell 15, 3709–3718 (2004).

62. P. Berardi et al., Both genome instability and replicative senescence stem from the shortest telomere in telomerase-negative cells. Nat Commun 17, (2026).

63. S. M. Dewhurst et al., Structural variant evolution after telomere crisis. Nat Commun 12, 2093 (2021).

64. R. C. Allsopp, C. B. Harley, Evidence for a critical telomere length in senescent human fibroblasts. Exp Cell Res 219, 130–136 (1995).

65. Y. Wu et al., Large-scale drug sensitivity, gene dependency, and proteogenomic analyses of telomere maintenance mechanisms in cancer cells. Nat Commun 16, 11337 (2025).

66. R. A. Dagg et al., Extensive Proliferation of Human Cancer Cells with Ever-Shorter Telomeres. Cell Rep 19, 2544–2556 (2017).

67. N. Viceconte et al., Highly Aggressive Metastatic Melanoma Cells Unable to Maintain Telomere Length. Cell Rep 19, 2529–2543 (2017).

68. J. K. Alder et al., Diagnostic utility of telomere length testing in a hospital-based setting. Proc Natl Acad Sci U S A 115, E2358–E2365 (2018).

69. S. Ackermann et al., A mechanistic classification of clinical phenotypes in neuroblastoma. Science 362, 1165–1170 (2018).

70. G. B. Morin, Recognition of a chromosome truncation site associated with alpha-thalassaemia by human telomerase. Nature 353, 454–456 (1991).

71. C. G. Kinzig, G. Zakusilo, K. K. Takai, L. R. Myler, T. de Lange, ATR blocks telomerase from converting DNA breaks into telomeres. Science 383, 763–770 (2024).

72. V. P. Schulz, V. A. Zakian, The saccharomyces PIF1 DNA helicase inhibits telomere elongation and de novo telomere formation. Cell 76, 145–155 (1994).

73. K. Myung, C. Chen, R. D. Kolodner, Multiple pathways cooperate in the suppression of genome instability in Saccharomyces cerevisiae. Nature 411, 1073–1076 (2001).

74. G. Cristofari, J. Lingner, Telomere length homeostasis requires that telomerase levels are limiting. EMBO J 25, 565–574 (2006).

75. N. T. Umbreit et al., Mechanisms generating cancer genome complexity from a single cell division error. Science 368, (2020).

76. Z. Zhang et al., Molecular insights into early malignant transition of hepatocellular carcinoma. Cancer Cell 44, 1012–1028 e1019 (2026).

77. F. K. Lorbeer, et al., Distinct senescence mechanisms restrain progression of dysplastic nevi. PNAS Nexus 3, pgae041 (2024).

78. I. T. P.-C. A. o. W. G. Consortium, Pan-cancer analysis of whole genomes. Nature 578, 82–93 (2020).

79. K. K. Takai, S. Hooper, S. Blackwood, R. Gandhi, T. de Lange, In vivo stoichiometry of shelterin components. J Biol Chem 285, 1457–1467 (2010).

80. S. Borowicz et al., The soft agar colony formation assay. J Vis Exp, e51998 (2014).

81. R. J. O’Sullivan, S. Kubicek, S. L. Schreiber, J. Karlseder, Reduced histone biosynthesis and chromatin changes arising from a damage signal at telomeres. Nat Struct Mol Biol 17, 1218–1225 (2010).

82. N. Arnoult et al., Regulation of DNA repair pathway choice in S and G2 phases by the NHEJ inhibitor CYREN. Nature 549, 548–552 (2017).

83. A. J. Cesare, M. T. Hayashi, L. Crabbe, J. Karlseder, The telomere deprotection response is functionally distinct from the genomic DNA damage response. Mol Cell 51, 141–155 (2013).

84. B. T. Letsolo, J. Rowson, D. M. Baird, Fusion of short telomeres in human cells is characterized by extensive deletion and microhomology, and can result in complex rearrangements. Nucleic Acids Res 38, 1841–1852 (2010).

85. N. Deimler, D. V. Ho, N. Paul, Z. Gill, P. Baumann, TARPON-A Telomere Analysis and Research Pipeline Optimized for Nanopore. PLoS Comput Biol 22, e1013915 (2026).

86. A. Dobin et al., STAR: ultrafast universal RNA-seq aligner. Bioinformatics 29, 15–21 (2013).

87. S. Heinz et al., Simple combinations of lineage-determining transcription factors prime cis-regulatory elements required for macrophage and B cell identities. Mol Cell 38, 576–589 (2010).

88. M. I. Love, W. Huber, S. Anders, Moderated estimation of fold change and dispersion for RNA-seq data with DESeq2. Genome Biol 15, 550 (2014).

89. J. M. Elizarraras et al., WebGestalt 2024: faster gene set analysis and new support for metabolomics and multi-omics. Nucleic Acids Res 52, W415–W421 (2024).

90. G. Yu, L. G. Wang, Y. Han, Q. Y. He, clusterProfiler: an R package for comparing biological themes among gene clusters. OMICS 16, 284–287 (2012).

91. Z. Gu, R. Eils, M. Schlesner, Complex heatmaps reveal patterns and correlations in multidimensional genomic data. Bioinformatics 32, 2847–2849 (2016).

92. H. Chen, P. C. Boutros, VennDiagram: a package for the generation of highly-customizable Venn and Euler diagrams in R. BMC Bioinformatics 12, 35 (2011).

93. Z. Gu, L. Gu, R. Eils, M. Schlesner, B. Brors, circlize Implements and enhances circular visualization in R. Bioinformatics 30, 2811–2812 (2014).

94. M. Diaz-Gay et al., Assigning mutational signatures to individual samples and individual somatic mutations with SigProfilerAssignment. Bioinformatics 39, (2023).

95. C. M. Grochowski et al., Inverted triplications formed by iterative template switches generate structural variant diversity at genomic disorder loci. Cell Genom 4, 100590 (2024).

96. J. R. Dixon et al., Integrative detection and analysis of structural variation in cancer genomes. Nat Genet 50, 1388–1398 (2018).

97. H. Li, Aligning sequence reads, clone sequences and assembly contigs with BWA-MEM. 2013 (10.48550/arXiv.1303.3997).

98. (Broad Institute, Broad Institute, GitHub repository, 2019).

99. N. Abdennur, L. A. Mirny, Cooler: scalable storage for Hi-C data and other genomically labeled arrays. Bioinformatics 36, 311–316 (2020).

100. M. D. Robinson, D. J. McCarthy, G. K. Smyth, edgeR: a Bioconductor package for differential expression analysis of digital gene expression data. Bioinformatics 26, 139–140 (2010).

101. A. Guarracino et al., Recombination between heterologous human acrocentric chromosomes. Nature 617, 335–343 (2023).

102. P. Danecek et al., Twelve years of SAMtools and BCFtools. Gigascience 10, (2021).

103. H. M. Amemiya, A. Kundaje, A. P. Boyle, The ENCODE Blacklist: Identification of Problematic Regions of the Genome. Sci Rep 9, 9354 (2019).

104. C. Zang et al., A clustering approach for identification of enriched domains from histone modification ChIP-Seq data. Bioinformatics 25, 1952–1958 (2009).

105. B. G. Stark R, DiffBind: differential binding analysis of ChIP-Seq peak data., (2011).

106. F. Ramirez et al., deepTools2: a next generation web server for deep-sequencing data analysis. Nucleic Acids Res 44, W160–165 (2016).

107. P. Hubel et al., A protein-interaction network of interferon-stimulated genes extends the innate immune system landscape. Nat Immunol 20, 493–502 (2019).

108. K. Chiba et al., Cancer-associated TERT promoter mutations abrogate telomerase silencing. Elife 4, (2015).

109. V. Valente et al., Selection of suitable housekeeping genes for expression analysis in glioblastoma using quantitative RT-PCR. BMC Mol Biol 10, 17 (2009).

